# Development of Small Molecule MEIS Inhibitors that modulate HSC activity

**DOI:** 10.1101/2020.02.12.946491

**Authors:** Raife Dilek Turan, Esra Albayrak, Merve Uslu, Pinar Siyah, Lamia Yazgi Alyazici, Batuhan Mert Kalkan, Galip Servet Aslan, Dogacan Yucel, Merve Aksoz, Emre Can Tuysuz, Neslihan Meric, Serdar Durdagi, Zafer Gulbas, Fatih Kocabas

## Abstract

*Meis1*, which belongs to TALE-type class of homeobox gene family, appeared as one of the key regulators of hematopoietic stem cell (HSC) self-renewal and a potential therapeutical target. However, small molecule inhibitors of MEIS1 remained unknown. This led us to develop inhibitors of MEIS1 that could modulate HSC activity. To this end, we have established a library of relevant homeobox family inhibitors and developed a high-throughput *in silico* screening strategy against homeodomain of MEIS proteins using the AutoDock Vina and PaDEL-ADV platform. We have screened over a million druggable small molecules *in silico* and selected putative MEIS inhibitors (MEISi) with no predicted cytotoxicity or cardiotoxicity. This was followed by *in vitro* validation of putative MEIS inhibitors using MEIS dependent luciferase reporter assays and analysis in the *ex vivo* HSC assays. We have shown that small molecules named MEISi-1 and MEISi-2 significantly inhibit MEIS-luciferase reporters *in vitro* and induce murine (LSKCD34^low^ cells) and human (CD34^+^, CD133^+^, and ALDH^hi^ cells) HSC self-renewal *ex vivo*. In addition, inhibition of MEIS proteins results in downregulation of *Meis1* and MEIS1 target gene expression including Hif-1α, Hif-2α and HSC quiescence modulators. MEIS inhibitors are effective *in vivo* as evident by induced HSC content in the murine bone marrow and downregulation of expression of MEIS target genes. These studies warrant identification of first-in-class MEIS inhibitors as potential pharmaceuticals to be utilized in modulation of HSC activity and bone marrow transplantation studies.

## 1. Introduction

*Meis1* is a member of TALE class of transcription factors ^1^. Through interaction domains in the N terminus, MEIS1 cooperates in transcription factors PBX1 and HOXA9 to transactivate target genes ^2,3^. MEIS2 and MEIS3 protein sequences demonstrate a high degree of similarity with MEIS1 ^4^. Meis1 was first described in leukemia mouse model and identified as a viral integration site (reviewed in ^5^). MEIS proteins are characterized by PBX interaction domains and a highly conserved homeodomain (HD). MEIS1 HD shares identical MEIS2 HD amino acid sequence. Studies to understand how MEIS1 HD interacts with DNA led to crystallization of MEIS1 HD with target DNA and identification of DNA sequence preferentially bound by MEIS proteins as “TGACAG” ^6–8^.

*Meis1* is highly expressed in the bone marrow ^2,9^. Lethality occurs in *Meis1* knockout mice at mid gestation (E14.5-15.5) with a number of hematopoietic, vascular and cardiac abnormalities ^10–12^. Conditional and tissue specific deletion of *Meis1* in bone marrow led to loss of HSC quiescence associated with expansion of HSC pool *in vivo* ^13^. *Meis1* has been shown to regulate HSC metabolism through transcriptional regulation of hypoxia factors including Hif1a and Hif2a ^13–16^. Deletion of *Meis1* or *Hif-1α* in HSCs leads to reduction in the cytoplasmic glycolysis and induction of mitochondrial phosphorylation. Intriguingly, studies showed a fundamental role of *Meis1* in neonatal cardiac regeneration. Increased *Meis1* expression was correlated with loss of neonatal cardiac regeneration, which is marked around day 7 ^17^. Cardiac specific deletion of *Meis1* in neonatal mice accelerated the cardiac regeneration post myocardial infarctions. These studies also demonstrated a transcriptional network that MEIS1 activates expression of a number of cyclin-dependent kinase inhibitors (CDKIs) in cardiomyocytes.

Numerous studies showed that MEIS1 is involved in pathways that regulate cell cycle, stem cell maintenance, redox state, cellular metabolism and carcinogenesis ^5^. These findings suggest that targeting MEIS1 could lead to development of new therapeutical approaches to treat numerous conditions where MEIS1 protein plays a pivotal role. However, lack of whole MEIS1 protein crystal structure, lack of MEIS specific *in vitro* drug screening tools, inefficient understanding of cellular response to *Meis1* loss of function, and issues regarding targeting a transcription factor by small molecules were core challenges for the development of small-molecule MEIS inhibitors. To this end, in the last decade, we have outlined how MEIS proteins could interact with target DNA, which is confirmed by published MEIS1 HD crystallization studies ^6,18^. In addition, we have developed several MEIS1-Luciferase reporter assays that could be applied to assess specificity of small molecule targeting MEIS1 activity ^13,16,17^. We have also outlined how loss of MEIS1 function, and how MEIS1 protein transcriptionally regulate *Hif-1α* and *Hif-2α* expression in hematopoietic compartment ^13^. Furthermore, we have showed that MEIS modulates expression of key CDKIs that could be used to assess efficacy of small molecule MEIS inhibitors by RT-PCR ^17^. Besides, further understanding of how other TALE family members interact with their respective target DNA, especially PBX1, PKNOX1, TGIF1 and TGIF2, provided us tools to develop small molecules inhibitors of MEIS1.

## 2. Materials and Methods

### 1. TALE family protein alignments

Protein sequences of TALE family members aligned to identify TALE family-conserved amino acids (aa). The aa sequences of each TALE family proteins were collected from NCBI. Multiple alignments were performed using the “constraint based multiple alignment tool” (COBALT) (NCBI). TALE family includes following; Myeloid Ecotropic Viral Integration Site 1 Homolog (MEIS1), MEIS2, MEIS3, Meis homeobox 3 pseudogene 1 (MEIS3P1), Meis homeobox 3 pseudogene 2 (MEIS3P2), PBX/knotted 1 homeobox (PKNOX1), PBX/knotted 1 homeobox 2 (PKNOX2), Transforming Growth Factor-Beta-Induced Factor 1 (TGIF1), TGFB induced factor homeobox 2 (TGIF2), TGFB induced factor homeobox 2 like, X-linked (TGIF2LX), TGFB induced factor homeobox 2 like, Y-linked (TGIF2LY), Pre-B-Cell Leukemia Transcription Factor 1 (PBX1), Pre-B-Cell Leukemia Homeobox 2 (PBX2), PBX homeobox 3 (PBX3), PBX homeobox 4 (PBX4), Iroquois Homeobox 1 (IRX1), IRX2, IRX3, IRX4, IRX5, IRX6, and Mohawk genes.

### 2. Generation of *in silico* small molecule library

*In silico* small molecule library used in this study was built by collecting small molecules from three different sources. First of all, the ZINC database drugs-now subset (http://zinc.docking.org/subsets/drugs-now) was used. The small molecules in the Drugs-now class are p.mwt <= 500 and p.mwt> = 150 and p.xlogp <= 5 and p.rb <= 7 and p.psa <150 and p.n_h_donors <= 5 and p.n_h_acceptors <= 10. In addition, the Sigma-Aldrich LOPAC1280 library was included in the study. Finally, known and potential inhibitors of homeobox gene family of proteins have been compiled from PubChem ^19,20^. Throughout compilation process, potential small molecules as homeobox protein family inhibitors were examined at PubChem for 314 genes. Then, homeobox inhibitor small molecule library was prepared in SDF format.

### 3. Molecular docking and cardiotoxicity predictions

The three-dimensional structure of the MEIS homeodomain was downloaded from PDB.org (PDB code: 3K2A) and then processed in the AutodockTools. Briefly, the water molecules were removed, the electrons in the atoms were checked and the crystal structure of the homeodomain was recorded in the form of pdbqt as we have done previously ^21,22^. Using AutoDockTools ^23^, pockets of conserved residues interacting with TGACAG MEIS binding DNA sequence were highlighted to determine the grid box locations with a 20-20-20°A search space. Docking studies were performed using AutoDock Vina 1.1.2 ^23^ and automated using PaDEL-ADV as we have done previously ^21,22^. Small molecules library including ZINC drugs-now supset (about 1 million compound) (**Supplementary File 1**), Sigma LOPAC1280 and Homeobox library of inhibitors were docked into MEIS homeodomain. The compounds with a binding energy of at least –6.9 kcal/mol were rescreened for the whole MEIS homeodomain surface with a 40-40-40°A search space, and determine the affinity to 4 other known TALE family proteins namely PBX(PDB:1DU6), PKNOX (PDB:1X2N), TGIF1 (PDB:2LK2), TGIF2LX (PDB:2DMN). Cardiotoxicity predictions were carried out by determining affinity to hERG channel as described previously ^24,24^.

### 4. Enrichment Analysis

Homeobox library inhibitors were expected to have lower free energies and expected enriched in terms of affinity towards MEIS homeobox protein. Thus, enrichment analysis was done by determining affinities of homeobox inhibitor library in comparison to a random ZINC drugs-now library and an unrelated SIGMA LOPAC1280 dataset. Affinities were interpreted in log scale and compared to enrichment in -6.9 kcal/mol, -7.0 kcal/mol, -7.1 kcal/mol and lower. Calculations are provided in **Supplementary File 2**.

### 5. Luciferase reporter assays

MEISi-1 (MW: 388.467) (Cas# 446306-43-0) and MEISi-2 (MW: 306.32) (Cas#2250156-71-7) are now available by Meinox Technologies (www.meinoxtech.com, Cat# MEISi1-25mg and Cat# MEISi2-25mg). Lyophilized MEISi-1 and MEISi-2 small molecules were dissolved in sterile dimethyl sulfoxide (DMSO, Calbiochem, 317275). Luciferase reporter assays were carried out as we have done previously ^13,14,16,17^. Briefly, p21-pGL2 (2 µg) (MEIS-p21-Luc reporter(^17^)) or Hif-pGL2 (MEIS-HIF-Luc Reporter or PBX-Luc reporter (^13,14,16^) was co-transfected with *Meis1* expression vector pCMVSPORT6-Meis1 or *Pbx1* pCMVSPORT6-Pbx1 (OpenBiosystems, 400 ng), pGL2 (filler) into HEK293 cells in six-well plates using poliethylenimine transfection reagent (SantaCruz, Cat.No. Sc-360988A). DMSO (control, 0.5%), MEISi-1 and 2 were tested in three different doses (0.1, 1 and 10 µM) in triplicates. After 48 hours of transfection, the cell lysate was processed for luciferase activity using the luciferase reporter system, according to the manufacturer’s instructions (Promega Dual-Glo, Cat.No. E2920). Luciferase measurements were done with Luminoskan Ascent Microplate (Thermo Lab System) and were calculated as relative light units normalized to transfection control / protein content.

### 6. Lineage negative cell isolation from mouse bone marrow

Mice were housed and provided by YUDETAM (Yeditepe University Experimental Animals Center). Animal studies were approved and performed in accordance with the relevant guidelines and regulations by the Institutional Animal Care and Use Committee of Yeditepe University (YUDHEK, decision numbers 418 and 446). Mouse whole bone marrow (WBM) was collected from the femurs and tibias of mice by flushing the bone marrow with insulin syringe and DPBS. WBM cells were filtered through 70-µM cell strainer (BD, Cat.No.352350). Isolated WBM cells were centrifuged and the supernatant was removed. The cells were resuspended into DPBS supplemented with two percent fetal bovine serum (FBS, v/v). Lineage negative (Lin-) cell fraction was obtained by elimination of Lineage + fraction based on a magnetic system following manufacturer’s recommendations. Briefly, WBM cells were blocked with Fc Blocker and stained with biotinylated**-**Lineage antibody cocktail according to the Mouse Hematopoietic Stem and Progenitor Cell Isolation Kit by BD (Catalog No. 560492). The stained cells were labelled with streptavidin magnetic particles and placed into BD IMagnet (BD, Cat.No. 552311). After fifteen minutes, labeled positive cells were attracted to the magnet and stick to the wall of falcon tubes as a pellet. Supernatant, which included Lin-fraction, was collected from the falcon tubes.

### 7. Flow cytometry analysis of murine HSPCs

Mouse bone marrow was extracted from the femurs and tibias of mice (YUDETAM) following euthanasia by flushing the bone marrow space with DPBS. Lin-cell fraction was isolated as described above. 30,000 Lin-cells/well were seeded in 96-well plates and treated with MEIS inhibitors for 7 days. Post 7 days of treatment, Lin-cells were collected and treated with Fc Blocker, then murine HSC surface markers including Lin-APC cocktail, Sca-1-PE-Cy7, c-Kit-PE, CD34-FITC according to manufactures recommendations (Mouse Hematopoietic Stem and Progenitor Cell Isolation Kit, BD, Catalog No. 560492). LSKCD34^low^ cells were analyzed by flow cytometer (BD FACS Calibur) as we have done previously^13–17,25–29^.

### 8. Lin-proliferation analysis by imaging

In order to determine the effect of MEISi-1 and MEISi-2 small molecules in HSPCs expansion, we utilized Cytell imaging platform (GE Healthcare). 10,000 Lin-cells were treated with DMSO, MEISi-1 and MEISi-2 (final concentrations; 0.1 µM, 1 µM, 10 µM) and were stained with Hoechst 33342 (Sigma Aldrich, Cat.No. 14533) at Day1 and Day 7. Photos covering a well of a 96-well plate were taken for each sample with Cytell imaging platform and Carl Zeiss Inverted Microscopy (ZEISS, Cat.No.849000464). The cell count per well was counted with Scion Image software.

### 9. Umbilical cord blood (UCB) and Peripheral Blood - Mononuclear Cell Isolation and Treatment

UCB mononuclear cells were isolated by Ficoll-Paque (Histopaque^TM^, Sigma, Cat.No. 10771) density gradient centrifugation (1.077 g/mL), and seeded in 96 well-plate in HSC medium at 10,000 cells per well for flow cytometry analysis as we have done previously ^14,16,25^. Briefly, 15 ml of cord blood was aliquoted into 50 ml falcon tubes (Isolab, CatNo. 078.02.004). Then, cord blood was diluted with 1:1 proportion of (v/v) DPBS. The diluted cord blood was added drop by drop into 1:1 proportion (v/v) of Ficoll-Paque. These mixtures were centrifuged (without brake). After centrifugation, cloudy interphase from obtained three different phases was collected, which includes mononuclear cells. Isolated mononuclear cells were washed with DPBS and centrifuged down (with brake). The isolated cells were seeded in the expansion medium which consists of Serum-Free Expansion Medium (StemSpan™ Serum-Free Expansion Medium (SFEM), Stemcell Technologies, Cat.No. 09650) supplemented with 1% PSA (10,000 units/ml penicillin and 10,000 µg/ml streptomycin and 25 µg/mL of Amphotericin B, Gibco, Cat.No.15240062) and human cytokine cocktail (StemSpan™ CC100, Stemcell Technologies, Cat.No. 02690). The seeded cells were treated with MEISi-1 and MEISi-2 small molecules at 0,1 µM, 1 µM and 10 µM concentrations. The cells used as positive control were treated with DMSO (0.5%) (Calbiochem, Cat.No.317275). The seeded cells were incubated at 37°C with 5% CO_2_ for 7 days.

Mobilized peripheral blood (mPB) mononuclear cells underwent Ficoll-Paque gradient, then 30,000 cells per well seeded in 96 well-plate (Corning CLS 3599) in human HSC medium as we have done previously ^14,16,25^. The human HSC medium consists of Serum-Free Expansion Medium (StemSpan™ Serum-Free Expansion Medium (SFEM), Stemcell Technologies, Cat.No. 09650) supplemented with 1% PSA (10,000 units/ml penicillin and 10,000 µg/ml streptomycin and 25 µg/mL of Amphotericin B (Gibco, Cat.No.15240062) and human cytokine cocktail (StemSpan™ CC100, Stemcell Technologies, Cat.No. 02690). The seeded cells were then treated with, DMSO (Control, 0.5%)(Calbiochem, Cat.No.317275) MEISi-1 and MEISi-2 small molecules at 0,1 µM, 1 µM and 10 µM concentrations. The seeded cells were incubated at 37°C with 5% CO_2_ until flow analysis. All human studies were approved by the Institutional Clinical Studies Ethical Committee of Yeditepe University (Decision numbers 547 and 548). All methods were performed in accordance with the relevant guidelines and regulations and appropriate informed consent were taken for human samples.

### 10. Flow cytometry analysis of human HSPCs

After 7 days of the treatment with the MEISi-1 and MEISi-2, the UCB- and PB-MNCs were labeled with PE-conjugated anti-human CD34 (Biolegend, Cat.No.343506), APC-conjugated anti-human CD133 (Miltenyibiotec, Order No.130-090-826) antibodies as we have done previously ^14,16,25^. In addition, UCB mononuclear cells were treated with Aldefluor reagent according to the manufacturer’s manual (ALDEFLUOR™ Kit, Stemcell Technologies, Cat.No. 01700). The expressions of CD34, CD133 surface markers and ALDH (Aldehyde Dehydrogenase) enzyme activity in the labeled cells were analyzed by flow cytometry (BD FACSCalibur™, Cell Analyzer) as we have done previously ^14,16,25^.

### 11. Colony forming assays

To perform colony-forming unit assay, the murine Lin-cells (30.000 cells/well) were treated with the 1 µM dose of MEISi-1 and MEISi-2. After 7 days expansion, the cells were harvested and counted. Then, the 20,000 cells/well were plated in methylcellulose-containing medium (MethoCult™ GF M3434, Stemcell Technologies, Cat.No. 03444) per well in 6-well plate, performed in triplicate. After 12 days, colonies were quantified under light microscopy as CFU-GEMM, CFU-G/M, BFU-E colonies as we have done previously^14,16,25,28,29^.

### 12. RT-PCR

Lin-cells isolated from mouse bone marrow were seeded on six well plate (Corning, CLS 3516) as one million cells/well. After four days of expansion, the cells were collected for RNA isolation. Total RNA was isolated by using Zymogen Quick RNA Mini Prep (ZymoResearch, Cat.No. R1054S) according to the manufacturer’s protocol. Isolated total RNA arranged as 2 µg total RNA and synthesized cDNA by following recommended protocol for SuperScript II Reverse Transcriptase Kit (Life Technology, 18080-051). Reaction was performed using with GoTaq qPCR Master Mix (Promega Cat.No6002) according to the manufacturer’s protocol. We have studied *Meis1*, *Meis1* target genes, CDKI and associated HDR genes post MEIS inhibitor treatments. Predesigned primers (**Table 1, Table 2, Table 3, and Table 4**) from PrimerBank (https://pga.mgh.harvard.edu/primerbank/) ^30–32^ were ordered from Sentegen. Real time PCR was performed by Light Cycler 96 (Roche Health Care Thermalcycler 96, Cat.No: 12953). GAPDH was used as a housekeeping control to normalize gene expression using the ΔΔCt method.

**Table 1.**
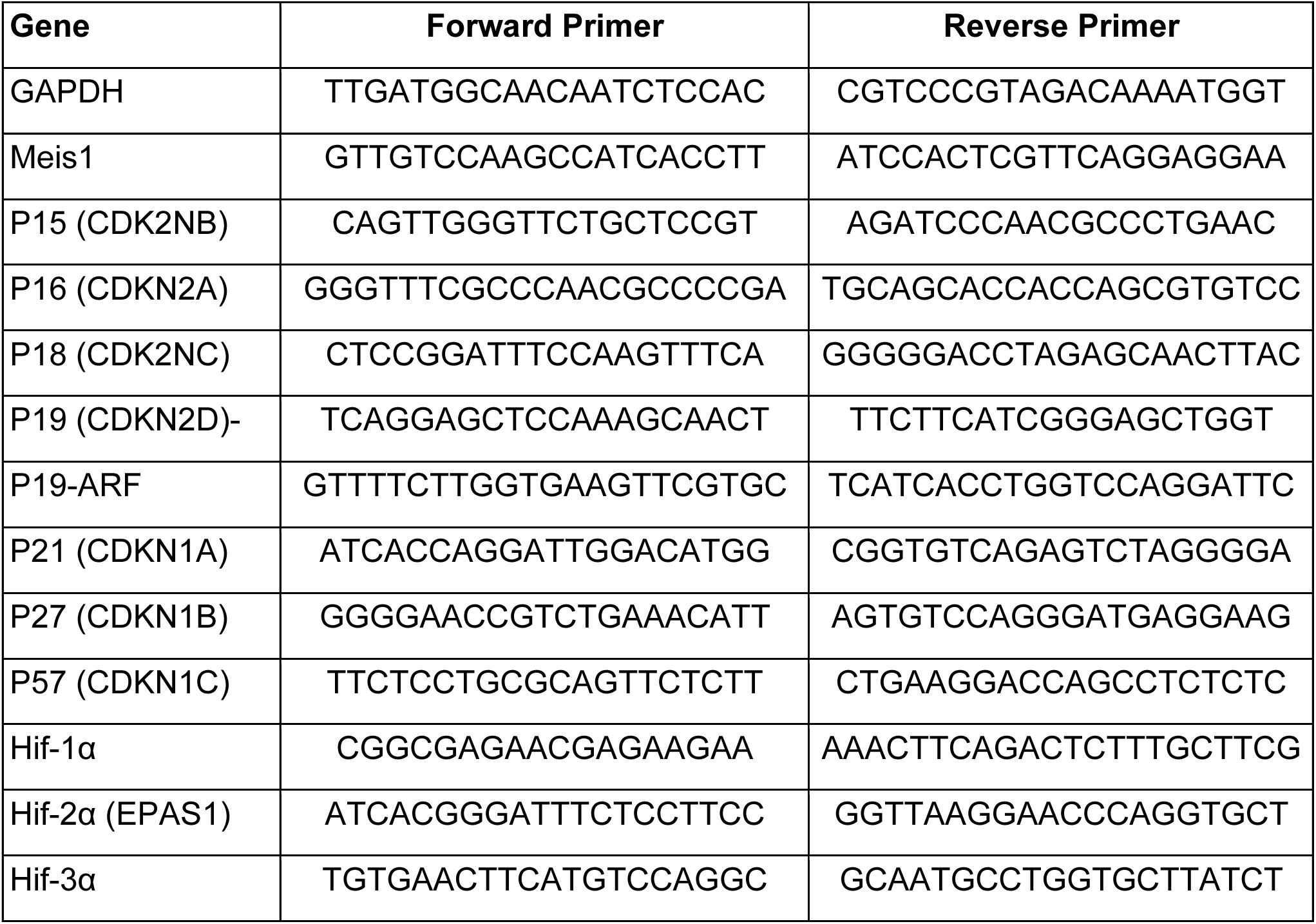
RT-PCR primer list for MEIS target and CDKI genes

**Table 2.**
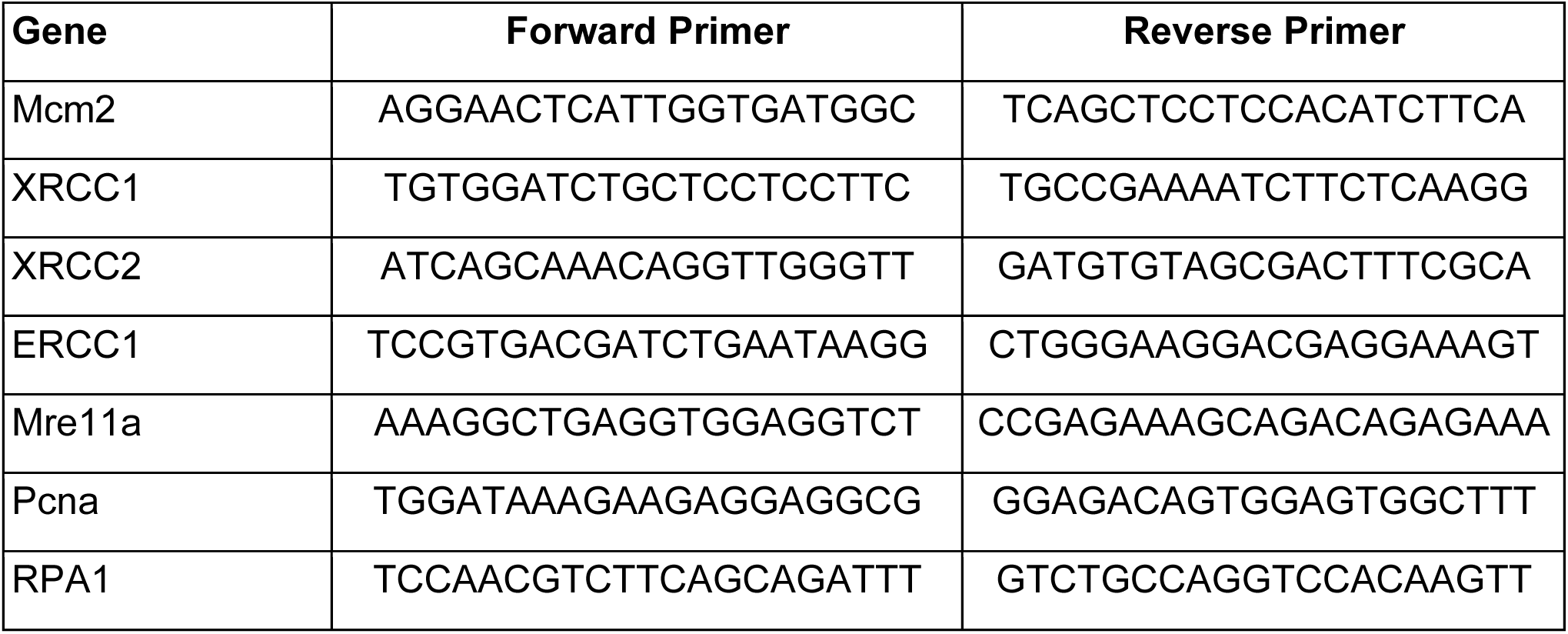
RT-PCR primer list for HDR and S-phase genes

**Table 3.**
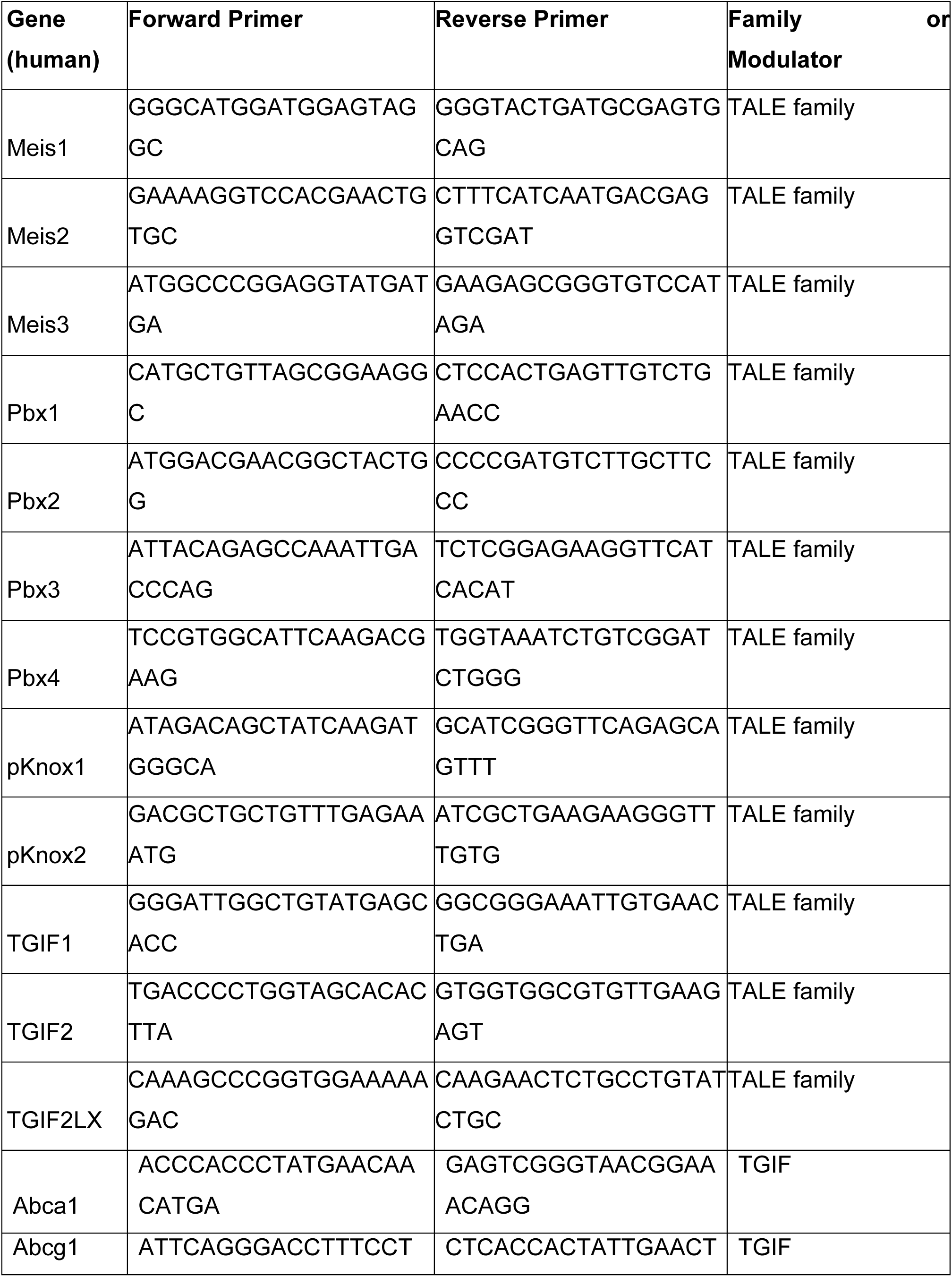
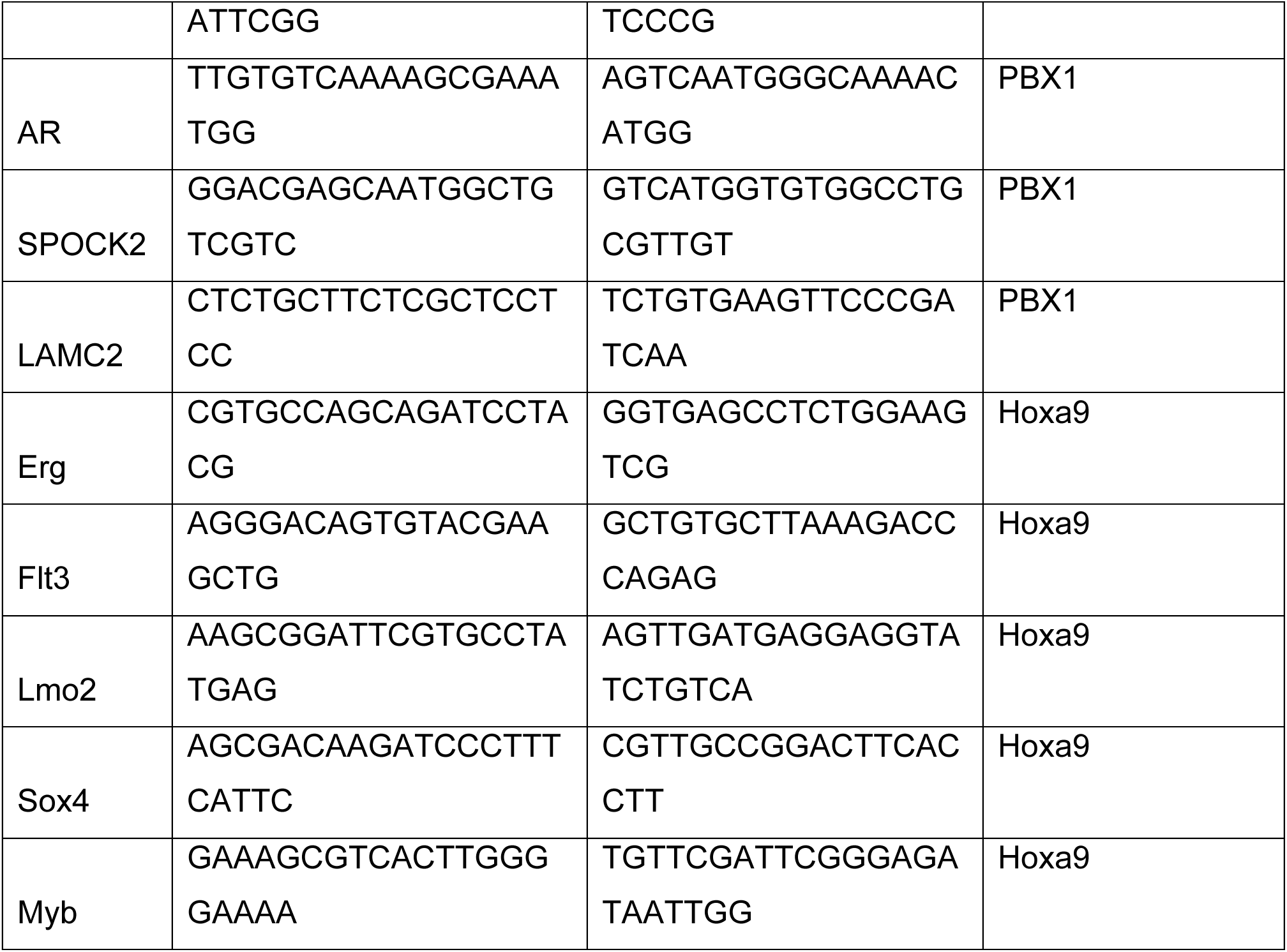
RT-PCR primer list for TALE Family, Pbx, Hoxa9 and TGIF target genes

**Table 4.**
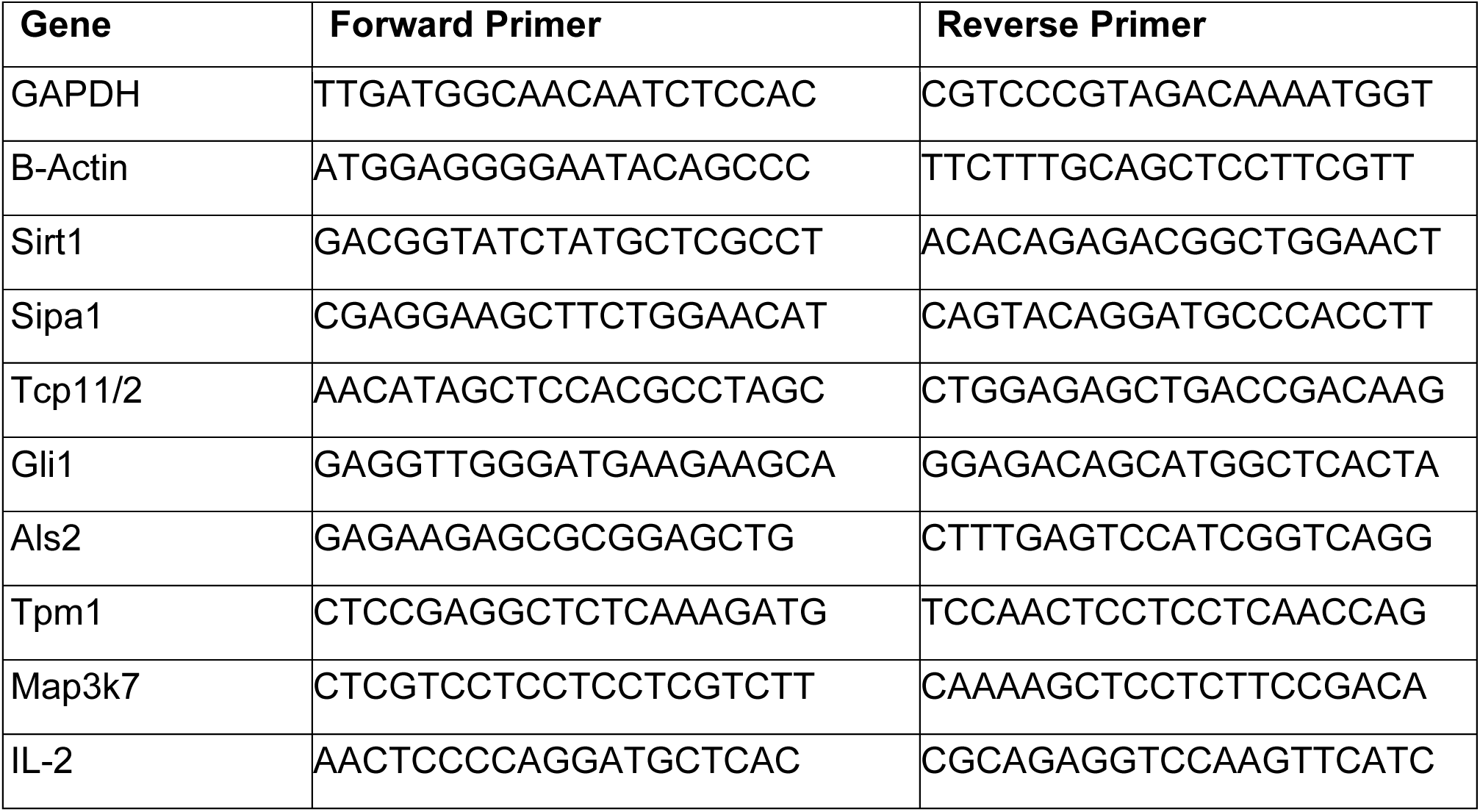
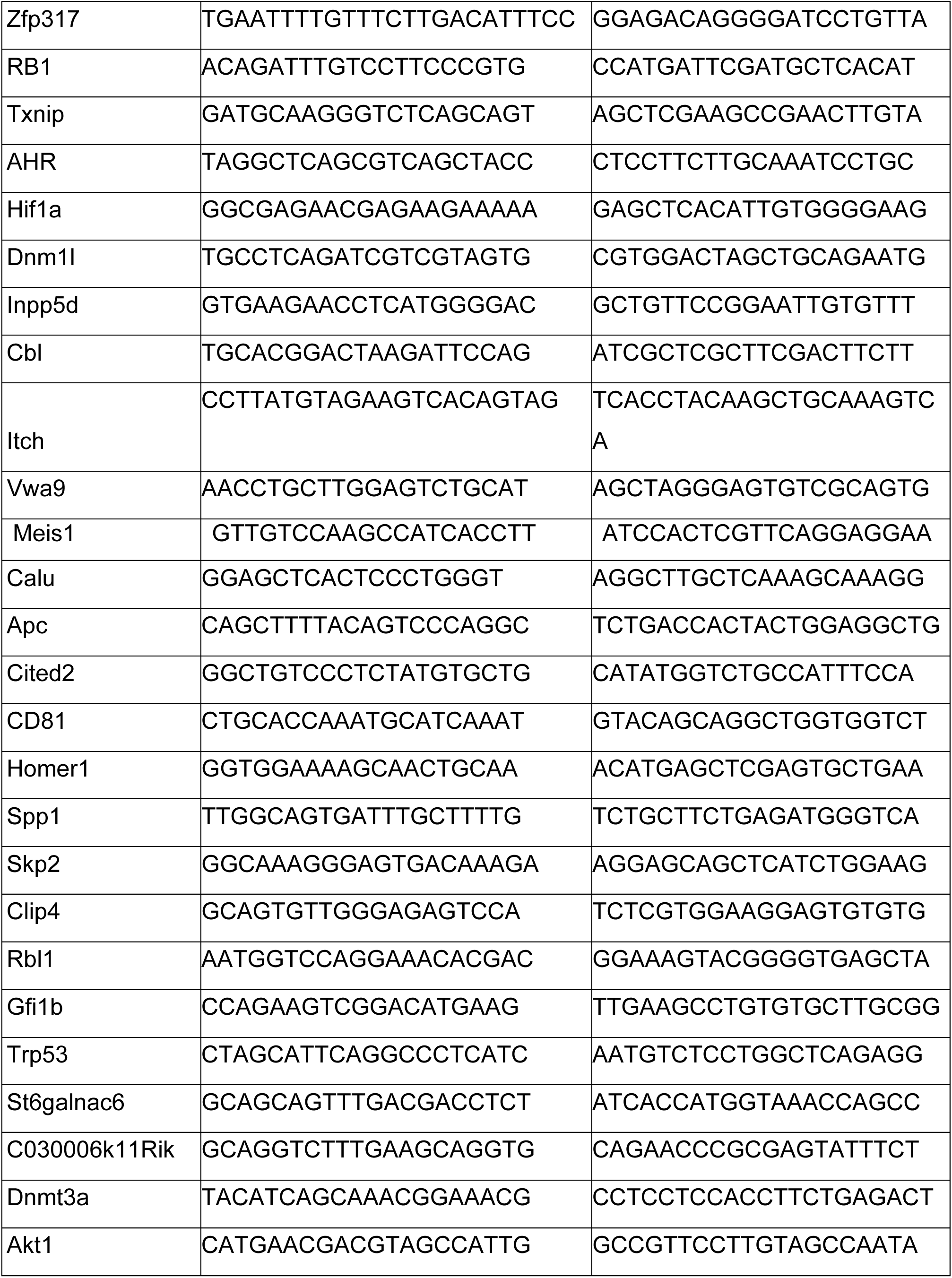
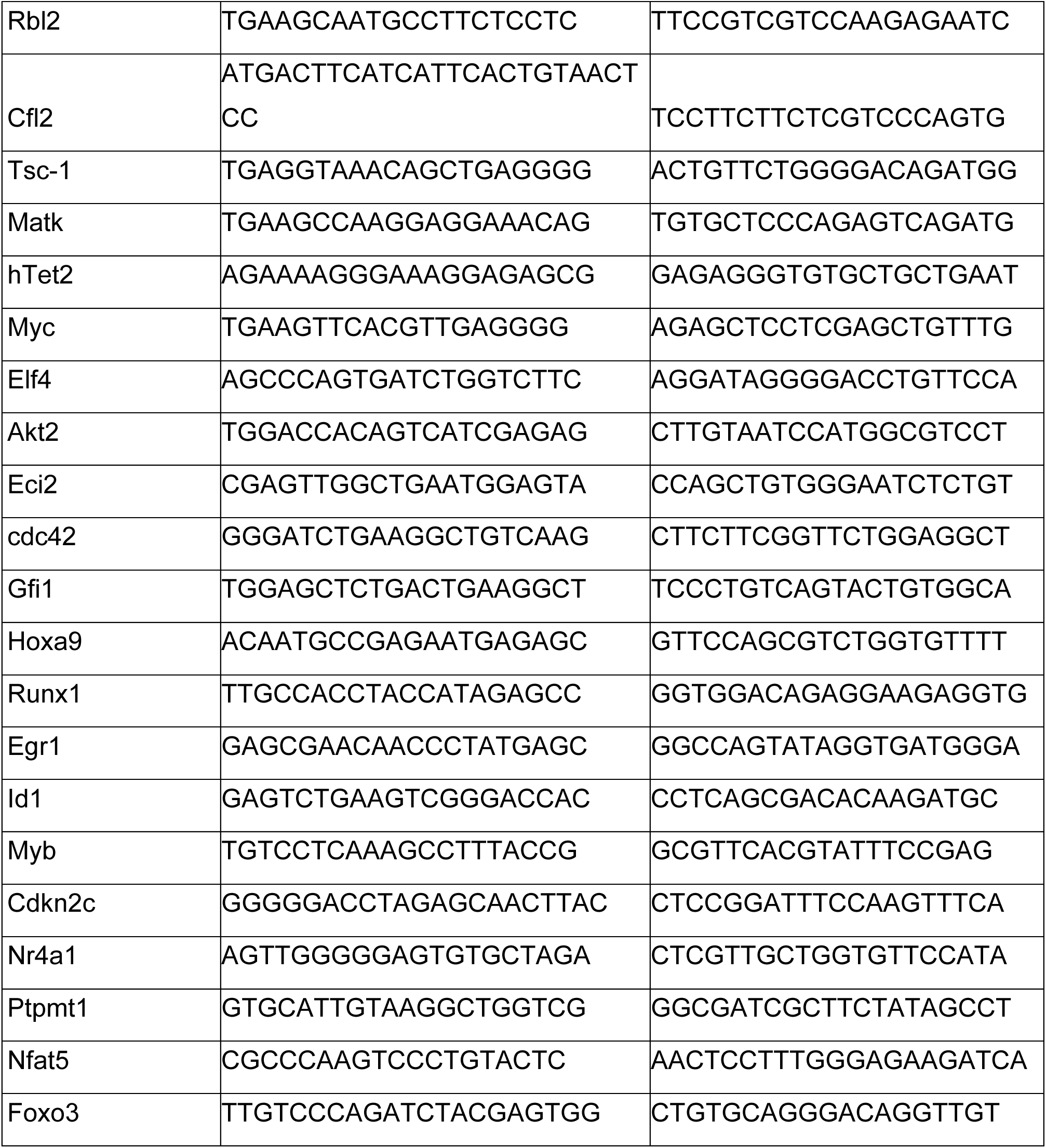
HSC gene pool primers

### 13. Mouse Bone marrow- and Human Adipose Derived-Mesenchymal Stem cell Isolation and Culture

Mouse whole bone marrow was flushed out from the femurs and tibias of Balb-C mice with insulin syringe and PBS as we have done previously ^25^. WBM cells were harvested as described above and seeded at a density of 30×10^6^ cells into T-75 flasks (Corning-Costar, Cat.No. CLS3290) in Dulbecco’s Modified Eagle’s Medium (DMEM Gibco, Cat.No.31885-023) supplemented with fifteen percent (v/v) FBS (Sigma Aldrich, USA, cat no. 12103C) and one percent (v/v) PSA (10,000 units/ml penicillin and 10,000 ug/ml streptomycin and 25 µg/mL of amphotericin B (Gibco, Cat.No.15240062). Mesenchymal stem cells (MSC) were treated with small molecules at passage number 3 (P3). BM-MSCs were seeded at a density of 10,000 cells/well in 96-well plate and treated with 0.1, 1 and 10 µM concentrations of MEISi-1 and MEISi-2 for WST1 cell proliferation assays.

Adipose tissue was collected by liposuction and processed within three to four hours. 150 ml adipose tissue was washed with equal volume of DPBS twice then tissue was placed into a 500 ml bottle, which comprise of equal volume of collagenase solution (0.2%, v/v) as we have done previously ^25^. The tissue was digested at 37°C for one to four hours by continuous shaking at 250 rpm. After the digestion, 10% FBS was added for inhibition of the collagenase activity. The digested tissue was centrifuged at 2000 rpm for ten min at room temperature. The supernatant that contains the collagenase solution and adipocytes was discarded and then, the pellet was resuspended in ammonium chloride solution (STEMCELL Technologies Cat. No:07850) for red blood cell lysis. The cell suspension was incubated at 37 °C for ten min by continuous shaking at 250 rpm. The cell suspension was washed centrifugation by using DPBS. Isolated cells were resuspended in DMEM, which were supplemented with 10% FBS and filtered through 100 µM cell strainer. The cells were seeded into T-75 flasks (Corning-Costar, Cat.No. CLS3290). Human AD-MSCs at passage two were used for small molecule treatments. After expansion of AD-MSCs, the cells were labeled with human MSC markers namely hCD73-APC, hCD90-FITC, hCD105-PerCP/Cy7, hCD45-PE according to the manufacturer’s protocol (not shown) and the flow cytometry analysis was performed to verify MSC origin of AD-MSCs. 10.000 cells per well were seeded in 96 well plates in 200 µl of MSC medium. The seeded cells were treated with MEISi-1 and 2 and analyzed for cell proliferation by WST1 assay post 3 days.

### 14. WST1 proliferation assays

Human umbilical vein endothelial cells (HUVECs, ATCC® CRL1730™) and primarily isolated BM-MSCs and AD-MSCs were studied for cell growth post treatment with MEIS inhibitors as we have done previously ^25,26^. Briefly, the cells were seeded in 96 well plates at a density of 5.000, 10.000 and 10.000 cells per well, respectively. MEISi-1, MEISi-2 and DMSO (Control, 0.5%) treatments were applied the next day of cell seeding. Post 3 days, 10µl / well cell proliferation reagent WST-1 (Boster, AR1159) added to cells. Cells were incubated with WST1 reagent for 2 hours in humidified cell culture incubator at 37°C and 5% CO_2_ in the dark. Absorbance was measured hourly at 450nm by Thermo Labsystem Multiskan Spektrum (Thermo, Cat.No.1500-176).

### 15. *In vivo* injections of MEIS inhibitors

In order to prove the influence of effective doses of MEISi determined by in vitro studies, BALB/C mice were used for in vivo studies. 10 mM concentration of MEISi-1 and MEISi-2 inhibitors were diluted with DPBS and final concentration adjusted as 10 µM. 100 µl of 1 µM MEISi-1, 10 µM MEISi-2, or 1% DMSO only were intraperitoneally injected into 4-6 week-old BALB/c mice at day 1, day 4 and day 7. At day 10, BALB/c mice sacrificed and whole bone marrow cells were isolated from femur and tibia for flow cytometry analysis and total RNA isolation. WBM cells were labeled with HSC LSKCD34^low^ surface markers, which are APC lineage cocktail, c-Kit (CD117) PE, Sca-1 PE-Cy7, CD34 FITC or HSC slam markers, which are APC lineage cocktail, c-Kit (CD117) PE, Sca-1 PE-Cy7, CD150 FITC and CD48 APC and analyzed with flow cytometry via FACS Calibur (BD Bioscience).

### 16. Repopulation analysis

*Ex vivo* expanded 500 HSCs (CD45.2+ homozygous) were transplanted into NOD/SCID mice homozygous for CD45.1 allele. Peripheral blood cells of recipient NOD/SCID CD45.1 mice were collected by retro-orbital bleeding, followed by lysis of red blood cells and staining with anti-CD45.2-FITC, anti-CD45.1-PE, anti-Thy1.2-PE (for T-lymphoid lineage), anti-B220-PE (for B-lymphoid lineage), anti-Mac-1-PE, or anti-Gr-1-PE (cells co-staining with anti-Mac-1 and anti-Gr-1 are deemed to be of the myeloid lineage) monoclonal antibodies (BD Pharmingen). The “percent repopulation” was determined based on the staining results of anti-CD45.2-FITC and anti-CD45.1-PE. Flow cytometry analysis was performed to confirm multilineage reconstitution.

### 17. Statistical analysis

Results are expressed as mean ± SEM. “2-tailed Student’s t test” was used to determine the level of significance. If the values had p˂0.05, results were considered statistically significant.

## 3. Results

### 3.1 Structure analysis, generation of screening library and generation of hits

The MEIS1 protein consists of Pbx interaction, homeodomain, and transcriptional activation domains. Pbx interaction domain, located at the N terminus, interacts with PBX1, PBX2, PBX3 proteins and is effective in its coactivation. The transactivation domain is located at the C-terminus. Homeodomain, which is 62 amino acid long, allows MEIS1 to bind specifically to DNA through recognition of TGACAG sequence. This specificity of MEIS1 HD makes it an ideal target for development of MEIS specific inhibitors. We first of all determined the key residues in the HD involved in DNA interaction. To this end, we blasted HDs of the TALE family of proteins to determine conserved amino acids (**Figure 1A**). In addition, we analyzed how MEIS HD interacts with DNA in crystallization studies (PDB # 5BNG). These studies allowed us to determine the key conserved amino acids in MEIS homeodomain that interact with target DNA. This is applied to other TALE family of proteins with similar HDs but different target DNA specificity. We have found that highly conserved F326, W327, R331, and R333 residues are found to be in proximity to and interact with target DNA. This is followed by generation of grid boxes around these four residues for each HDs of MEIS1, PBX1, TGIF1, TGIF2X, and PKNOX proteins (**Figure 1B**). We have also generated a grid box to perform whole surface docking studies to eliminate non-specific hits in downstream applications (**Figure 1B**).

**Figure 1.**
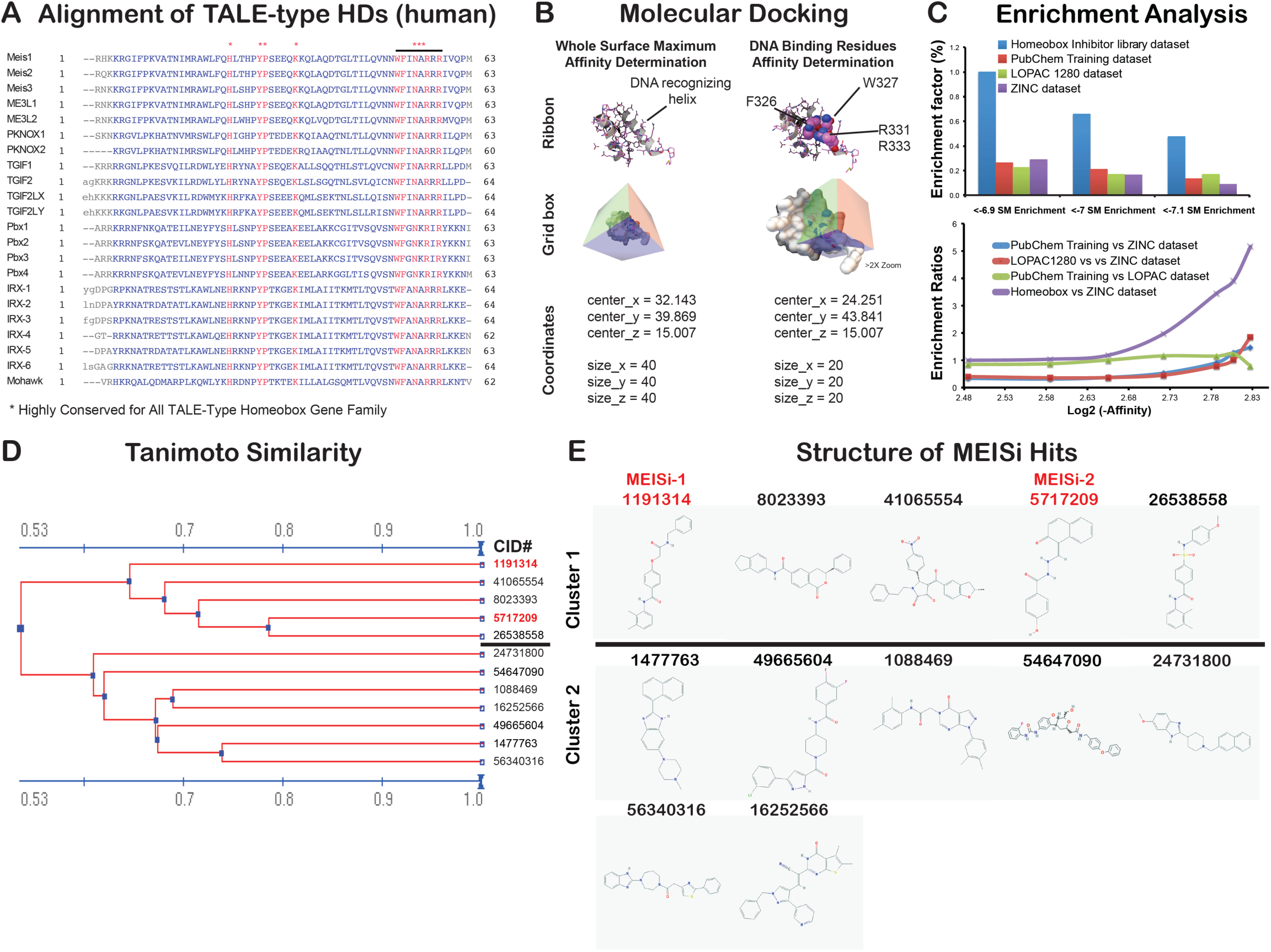
Development of MEIS-HD inhibitors. **A) Alignment of TALE type HDs.** The amino acid sequences of each TALE family proteins collected from NCBI. Multiple alignments were done using the “constraint based multiple alignment tool” (COBALT). Highly conserved residues are marked with *. Note that there are several 100% conserved residues across TALE family of proteins. **B) Molecular docking of MEIS homeodomain.** MEIS homeodomain was docked with small molecules in two grid boxes, whole surface included grid box and DNA binding residues grid box. Whole surface search box is performed to determine and eliminate small molecules non-specifically binds to homeodomain versus DNA binding residues. Dimensions in the grid box are provided in angstroms. **C) Enrichment analysis.** Percent enrichment and enrichment factor ratio analysis of hits in homeobox library of small molecules docked into MEIS HD compared to random (ZINC drugs-now set) or unrelated (Lopac1280 set, or PubChem training set) libraries. **D) Analysis of MEISi hit small molecules.** MEIS hit molecules analyzed for structural similarity based on substructure key-based 2D Tanimoto similarity. **E)** Two clusters of MEISi hit molecules were identified. Structures and CID numbers are provided.

We have performed *in silico* screening against MEIS HD in a three major steps. These included 1) generation of small molecule library, 2) automated docking with Autodock Vina and Padel ADV platform, and 3) elimination of non-specific hits of MEIS HD by docking into other HDs in the TALE family (**Figure S1**). Small molecule library largely included ZINC database All-purchasable-drugs-now subset (>1M compounds), LOPAC and MyriaScreen compounds (11.280 structures), PubChem tranining set (10.000 compounds) and in-house curated library of Homeobox related small molecule inhibitors reported to target homeobox family of proteins. In-house library of homeobox inhibitors were curated from PubChem database. To do this, we have collected small molecules targeting each of the 314 homeobox proteins in PubChem. We have downloaded the appropriate formats (SDF format) of these molecules in three-dimensional format for *in silico* screening studies. We determined that majority of homeobox inhibitors were associated with a dozen of homeobox proteins. In-house library of small molecules also allowed us to analyze the enrichment and targeting specificity to HD by comparing to affinities obtained in unrelated small molecule libraries (**Figure 1C**). Enrichment analysis demonstrated that screening strategy could successfully identified small molecules targeting homeodomain and it has more than 5 fold enrichment compared to unrelated Lopac1280, ZINC, or PubChem training sets (**Figure 1C**).

*In silico* docking and screening studies were done for generated grid boxes (grid box information provided in **Figure 1** and **Figure S2**) as we have done previously ^21,22^. We have completed docking of over 1M small molecules into MEIS HD (**Supplementary File 1**). Selected hits were also docked into whole protein, and other HDs of TALE proteins including PBX, PKNOX, TGIF1 and TGIF2LX. Binding affinity difference between MEIS HD vs other TALE HD were determined. Differences higher than 0.5 kcal/mol were selected. In addition, we have performed hERG channel dockings to eliminate potential cardiotoxic hits. These studies allowed us to narrow down to identify 12 putative and specific small molecule inhibitors of MEIS (**Table 5, Supplementary File 3**). Structural clustering of hits allowed us to determine similar hits (**Figure 1D**) and showed that these hits could be clustered into two (**Figure 1E**). Intriguingly, all hit compounds had polar molecules, especially oxygen in the central pocket. This is thought to relate to mimicking the negative nature of target DNA to competitively block interaction of MEIS HD with DNA.

**Table 5.** Selected MEIS specific hit small molecules.

| Ligand ID |  | Affinities (AutoDock Vina, kcal/mol) |  |  |  |  |  | Glide XP |
| --- | --- | --- | --- | --- | --- | --- | --- | --- |
| CID # | ZINC ID | MEIS<br>(PDB:3K2A) | PBX<br>(PDB:1DU6) | PKNOX<br>(PDB:1X2N) | TGIF1<br>(PDB:2LK2) | TGIF2LX<br>(PDB:2DMN) | ΔMEIS vs<br>other TALEs | hERG<br>channel |
| 1477763 | ZINC01390248 | -7.4 | -6.4 | -6.9 | -6.7 | -6.1 | 0.8 | -7.2 |
| 41065554 | ZINC09425201 | -7.3 | -5.6 | -6.2 | -5.2 | -5.2 | 1.8 | -7.2 |
| 49665604 | ZINC49406820 | -7.3 | -6.6 | -6.3 | -6.9 | -6.5 | 0.7 | -7.6 |
| 16252566 | ZINC16051182 | -7.3 | -6.7 | -6.9 | -7 | -6.5 | 0.5 | -6.3 |
| 1191314 | MEISI-1 | -7.2 | -6.2 | -6.4 | -6.2 | -6.3 | 0.9 | -7.1 |
| 56340316 | ZINC71853541 | -7.2 | -6.5 | -6.8 | -6.8 | -6.3 | 0.6 | -7.5 |
| 26538558 | ZINC15354059 | -7.1 | -5.7 | -6.6 | -6.1 | -5.7 | 1.1 | -7.5 |
| 24731800 | ZINC24880387 | -7.1 | -6.3 | -6.9 | -6.5 | -6.7 | 0.5 | -7.0 |
| 1088469 | ZINC00807131 | -7.1 | -6.5 | -7 | -6.6 | -6 | 0.6 | -7.6 |
| 8023393 | ZINC06647669 | -7.1 | -6.4 | -6.7 | -6.1 | -7 | 0.6 | -7.7 |
| 24731800 | ZINC24880387 | -7.1 | -6.3 | -6.9 | -6.5 | -6.7 | 0.5 | -7.0 |
| 5717209 | MEISI-2 | -6.9 | -6.5 | -6.5 | -6 | -5.9 | 0.7 | -6.9 |

Hit small molecules provided in chemical identification number (CID, PubChem), ZINC ID with affinities calculated against MEIS, PBX, PKNOX, TGIF1, and TGIF2LX with AutoDockVina. Glide XP was used to calculate affinities against hERG channel to predict any cytotoxicity. Note that hits were selected based on lower binding energy compared to other TALE family of proteins.

### 3.2 Selected compounds inhibited MEIS-Luc reporter

We have previously developed luciferase reporters that could be activated with introduction of MEIS1 protein ^13,14,16,17^. Once MEIS1 protein is expressed, it binds to conserved TGACAG motif located in the promoter region of MEIS-Luc reporter and transactivates expression of luciferase reporter (**Figure 2A**). We have tested selected compounds in MEIS-luc reporter assay following transfection of MEIS-luc, and pCMV-SPORT6-Meis1, and internal control pCMV-LacZ plasmids into HEK293T cells. After transfection, we have applied selected compounds in 0.1 µM, 1 µM, and 10 µM concentrations. Post 48 hours, we have measured luciferase activity and normalized to transfection control. MEIS-Luc reporter activity was significantly reduced with MEIS inhibitor 1 (MEISi-1) and MEIS inhibitor 2 (MEISi-2) at 0.1 µM concentration (**Figure 2B-C**) when tested against two different MEIS-Luciferase reporters. MEISi-1 and MEISi-2 also demonstrated a dose dependent activity (**Figure S3**). We have achieved up to 90% inhibition of MEIS-Luc activity with MEIS-1 and 2. Other tested three MEISi-1 and MEISi-2 related compounds from the in silico screening hits showed no inhibition (**Figure S4**). Besides, we have tested MEISi-1 and MEISi-2 in PBX-Luc reporter that we have previously characterized ^14^. We have shown that MEISi-2 does not significantly inhibit the PBX-Luc-reporter while MEISi-1 could inhibit only at higher doses (**Figure S5**).

**Figure 2.**
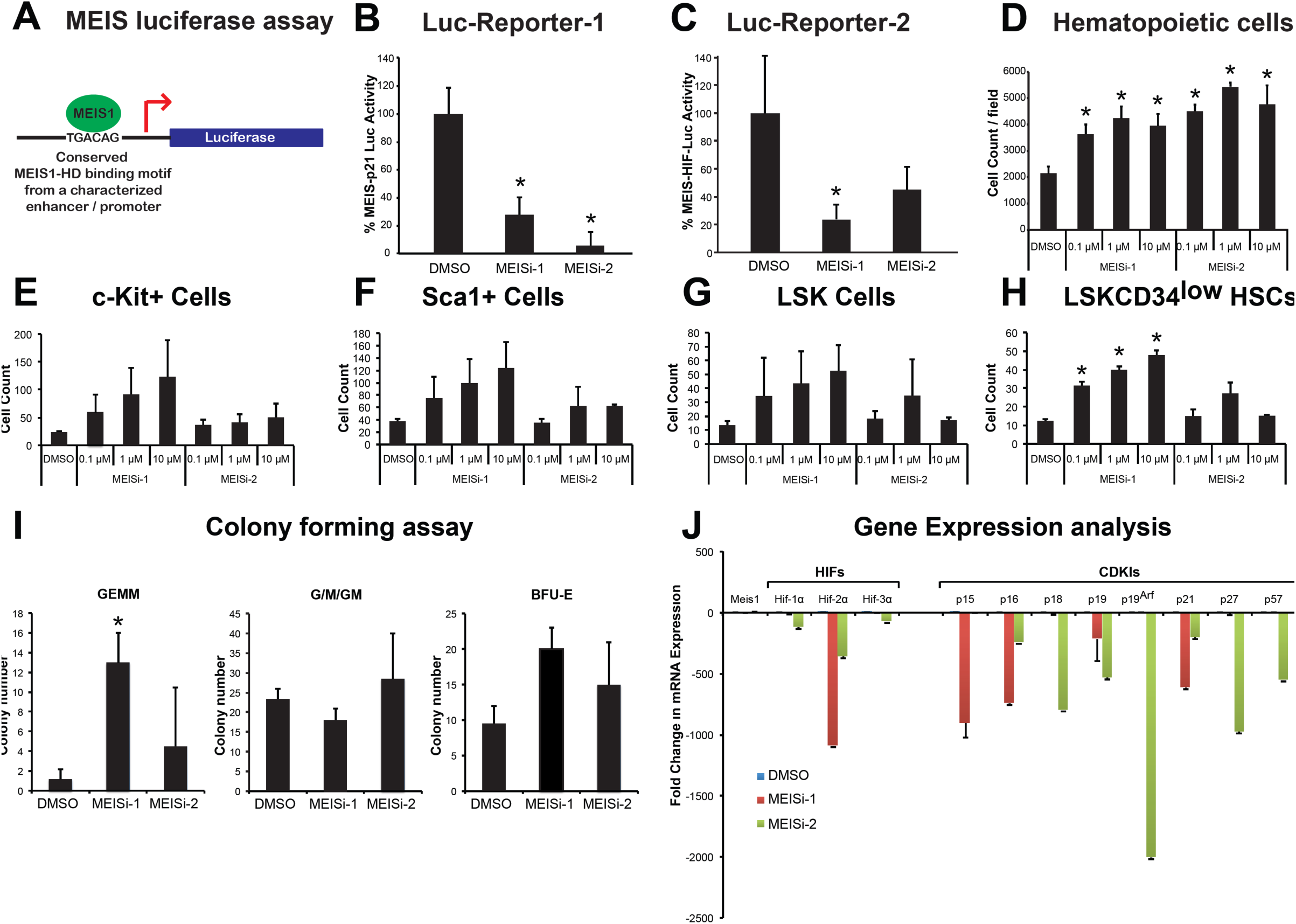
**A) MEIS luciferase reporter assay.** Schematic showing luciferase reporter with MEIS binding motif TGACAG in the regulatory regions that is used to test activity of Meis1 protein. **B) MEIS-Luc-Reporter 1.** Two small molecules named MEISi-1 and MEISi-2 demonstrated inhibition of MEIS-p21-luciferase reporter (luciferase reporter with well characterized MEIS1 binding motif from *p21* regulatory region) up to 90% at 0.1 µM concentration. **C) MEIS-Luc-Reporter 2.** MEISi-1 and MEISi-2 demonstrated inhibition of MEIS-HIF-luciferase reporter (luciferase reporter with well characterized MEIS1 binding motif from Hif-1α enhancer region). **D) Effect of MEISi treatments on the Lin-cells ex vivo.** Lin-cells were isolated and treated with corresponding MEIS inhibitors and doses. Post 7 days of treatment, cells were stained with Hoechst 33342 and counted using automated cell imaging platform. **Hematopoietic stem and progenitor cell expansion post MEISi treatments.** Lin-cells were isolated and treated with corresponding MEIS inhibitors and doses. Post 7 days of treatment, HSCs were stained with corresponding surface antigens **E) c-Kit^+^**, **F) Sca1^+^**, **G) LSK** and **H) LSKCD34^low^** and determined HSC content by flow cytometry. **I) Colony forming assay.** Lin-cells were treated with MEIS inhibitors for seven days. Then, methocult based CFU assays were performed. Types of colonies formed post 12 days were quantified and illustrated as CFU-GEMM, CFU-G/M/GM and BFU-E colonies. **J) Expression of MEIS target genes, HIFs and CDKIs post MEISi treatments in Lin-Cells.** Lin-cells were treated *in vitro* with MEIS inhibitors and collected RNA post 3 days of treatment for analysis of gene expression. Note that MEIS1 is known to transcriptionally regulate expression of *Hif-1α, Hif-2α, p15, p19^ARF^* and *p21*. n=3, *p<0.05

### 3.3 Selected compounds induced MEIS dependent hematopoietic stem cell maintenance and self-renewal

We have previously shown that *Meis1* knockout in HSC compartment leads to up to 5 fold expansion of HSC pool *in vivo*, albeit with increased apoptosis ^13^. We hypothesized that temporal inhibition of hematopoietic cells could allow *ex vivo* HSC expansion without apoptosis issue. Thus, we have isolated Lin-cells, which includes about 1% HSCs, and treated with MEISi-1 and MEISi-2. We have found that MEIS inhibitors induce hematopoietic cell count in a dose dependent manner (**Figure 2D**). In addition, analysis of HSC content by flow cytometer showed that MEIS both MEISi-1 and MEISi-2 induce c-Kit+ progenitor (**Figure 2E**), Sca1+ progenitor (**Figure 2F**), LSK HSPCs (**Figure 2G**), and LSKCD34low HSC count (**Figure 2H**) *ex vivo* in a dose dependent manner. MEISi-1 is more potent compared to MEISi-2 in the expansion of HSCs. This is further verified with CFU assays, which is an *in vitro* surrogate self-renewal assay to assess proper expansion HSCs. Here we show that MEISi-1 and MEISi-2 treatments induce number of mix colonies derived from HSCs (GEMM colonies, **Figure 2I**) up to 5 fold compared to DMSO control. Numbers of CFU-G/M/GM colonies were similar. Numbers of BFU-E colonies were slightly higher, albeit not significant, with MEISi-1 treatments. This findings suggest that MEIS inhibitors not only expand HSCs ex vivo but also maintains their self-renewal in the duration of treatments.

### 3.4 MEIS inhibitors downregulated MEIS target gene expression

We have previously outlined a number of molecular pathways under regulation of MEIS homeodomain ^5,13–17,27,28,33–35^. We have shown that MEIS1 transcriptionally activate *Hif-1α* and *Hif-2α* expression HSCs. Besides, we have recently shown that MEIS blocks neonatal cardiac growth by transcriptional activation of a number of CDKIs, more specifically *p15*, *p19^ARF^*, and *p21*. Here we tested how MEIS inhibitors we have identified affected the gene expression of MEIS1 target genes. We have found that MEISi-1 and MEISi-2 treatments downregulated expression of *Hif-1α* and *Hif-2α* in hematopoietic cells as we have expected (**Figure 2J**). In addition, expression of a number of CDKIs was largely downregulated with MEISi-1 or MEISi-2 treatments. Further analysis of *Meis, Pbx, Hoxa9, TGIF* and *pKnox* genes and their known targets by RT-PCR showed that, while we achieve significant downregulation of *Meis1* and *Meis2* expression and their target genes, we did not observe any significant change in expression profile of majority of related genes analyzed (**Figure S6**). These findings demonstrated the specificity and functional inhibition of MEIS HD by MEIS inhibitors that we identified in our *in silico* screening. In addition, we have determined gene expression by PCR array post MEISi-1 to determine how MEIS inhibitors affect expression of HSC related genes. We have found that expression of *Sirt1, Sipa1, Tcp11/2*, and *Gli1* were upregulated post MEISi-1 treatments (**Figure S7**). On the other hand, expression of *Foxo3, Nfat5*, and *Ptpmt1* were downregulated post MEISi-1 treatments. These findings suggest that MEIS inhibitors may modulate key HSC stem quiescence genes.

### 3.5 MEIS inhibitors are affective in targeting MEIS protein *in vivo* and modulate HSC pool

We next moved to address the functionality of MEIS inhibitors *in vivo*. We have known that HSC specific Meis1 deletion leads to a significant induction of HSC pool *in vivo*. We injected MEIS inhibitors into wild type mice by I.P. for 3 successive days, and collected bone marrow for HSC analysis (**Figure 3A**). We have found that both MEISi-1 and MEISi-2 functionally inhibit MEIS *in vivo*, thus they induce c-Kit+ cell (**Figure 3B**), Sca1+ cell (**Figure 3C**), CD150+ cell (**Figure 3D**), LSK HSPCs (**Figure 3E**), LSKCD34low HSC content (**Figure 3F**) and LSKCD150+CD48-HSC content (**Figure 3G**).

**Figure 3:**
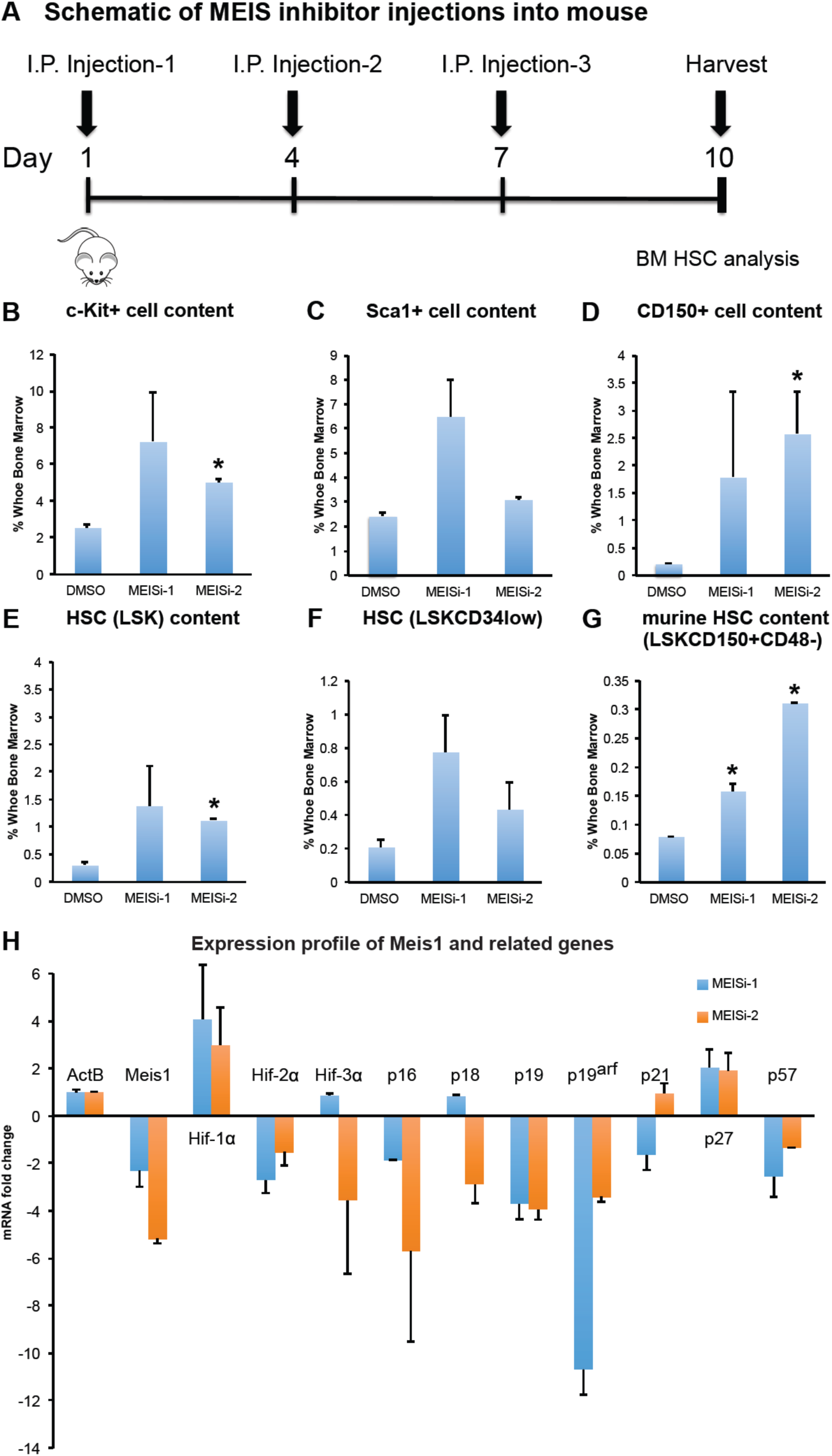
Phenotypical HSC antigen analysis post MEISi injections. **A)** Schematic showing the intraperitoneal injections of MEIS inhibitors and BM analysis at day 10. *In vivo* analysis of HSC compartment post MEISi treatments was carried out by analysis of **B)** c-Kit^+^, **C)** Sca-1^+^ cell content, **D)** CD150^+^ cell content, **E)** LSK cell content, **F)** LSKCD34^low^ cell content and **G)** LSKCD48^-^CD150^+^ HSC content in the whole bone marrow following injection of MEISi-1, MEISi-2 and DMSO control. **H)** Expression profile of Meis1 and related target genes post MEISi injections in the whole bone marrow cells. n=3, *p<0.05

We also performed gene expression profiling of bone marrow in MEISi-1 and MEISi-2 injected animals. We have verified that both MEISi-1 and MEISi-2 treatments led to downregulation of *Meis1* expression, *Hif-2α*, and key CDKI gene expression including *p16*, *p19, p19ARF* (**Figure 3H**). Intriguingly, we have not seen downregulation of *Hif-1α* as we have expected but we have observed downregulation of Hif-2α expression in the bone marrow post MEISi-1 and MEISi-2 treatments. These finding, overall, demonstrate that MEISi-1 and MEISi-2 could functionally used to target MEIS protein and associated pathways *in vivo*.

### 3.6 MEISi allowed human UBC HSCs and BM HSC maintenance and expansion *ex vivo*

HSC expansion technologies become more important since the development of tools to edit inherited mutations in HSCs. These procedures involve single cell selection and expansion of HSCs to large numbers to achieve efficient engraftment abilities in treatment of patients especially suffering from inherited forms of anemia. To this end, we assessed if newly developed MEIS inhibitors could be utilized in UCB HSC or human mobilized peripheral blood (mPB) HSC expansion procedures.

Isolated UCB mononuclear cells were treated with three different doses (final concentrations; 10 µM, 1 µM, 0.1 µM) of MEISi-1 and MEISi-2. After the seven days of the treatment, the cells were characterized for the human HSC markers (**Figure S8**). We have found that MEISi-1 and MEISi-2 treatments significantly increase human UCB hematopoietic cell count up to 5 fold compared to DMSO (**Figure 4A**). In addition, CD34+ (**Figure 4B**), CD133+ (**Figure 4C**), and ALDH^br^ (**Figure 4D**). UCB HSPC cell content was increased up to 4 fold with MEISi-1 and MEISi-2 treatments.

**Figure 4.**
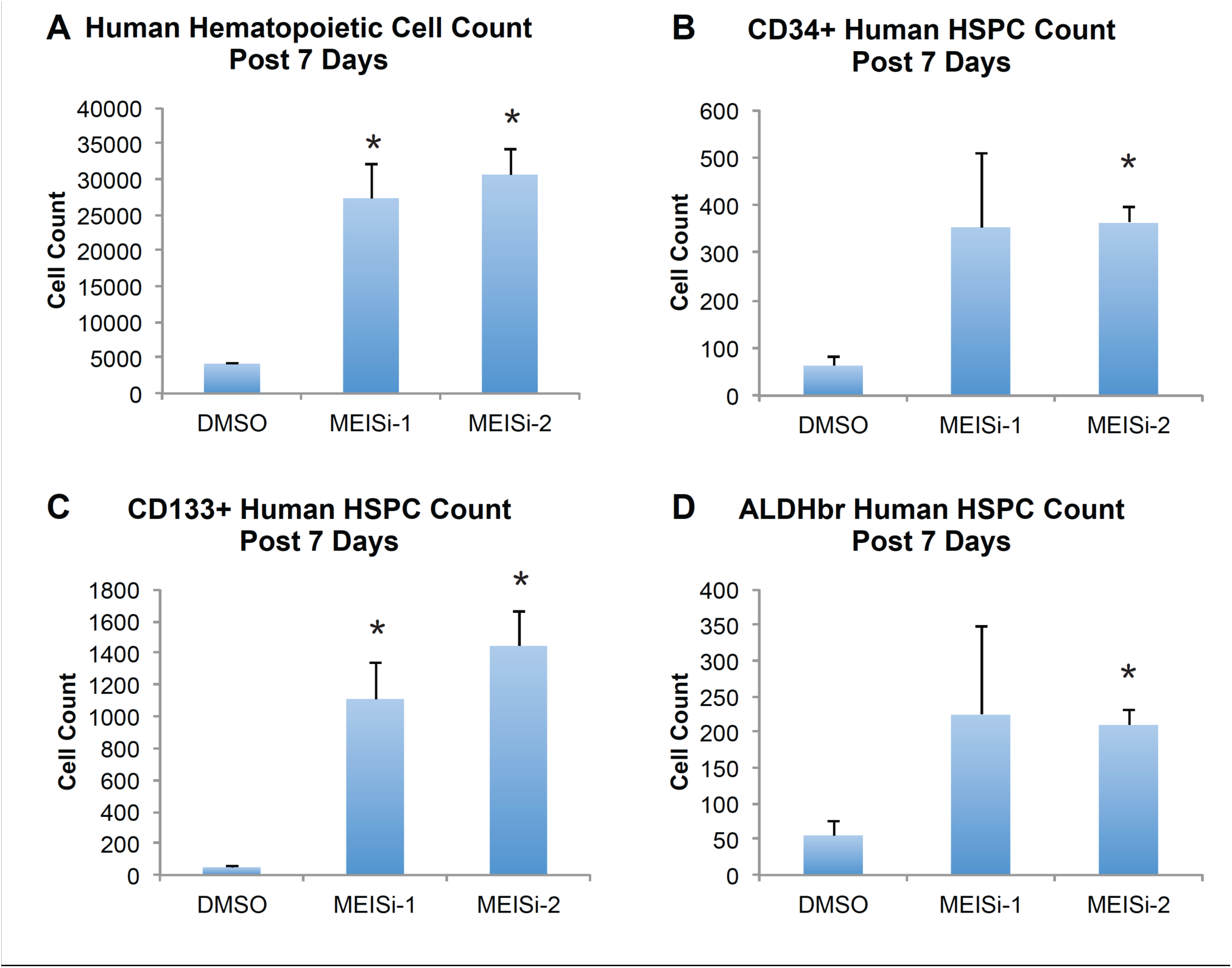
Human UCB hematopoietic cell expansion post MEISi treatments. **A)** Human Hematopoietic cell count post 7 days of MEISi treatments, **B)** CD34^+^ human HSPC count post 7 days, **C)** CD133^+^ human HSPC count post 7 days, **D)** ALDH^br^ human HSPC count post 7 days. n=3. *p <0.05.

We have also obtained human mPB HSCs and treated with MEISi-1 and MEISi-2. We achieved induction of CD34+ (**Figure 5A**), CD133+ (**Figure 5B**), CD90+ (**Figure 5C**), CD34+CD38- (**Figure 5D**), CD34+CD133+CD90+ (**Figure 5E**), and CD34+CD133+CD90+CD38- (**Figure 5F**) human mPM HSCs at least 2 fold with MEISi-1 treatments. Intriguingly, we only observed induction of CD133+ population with MEISi-2 treatment (**Figure 5B**).

**Figure 5.**
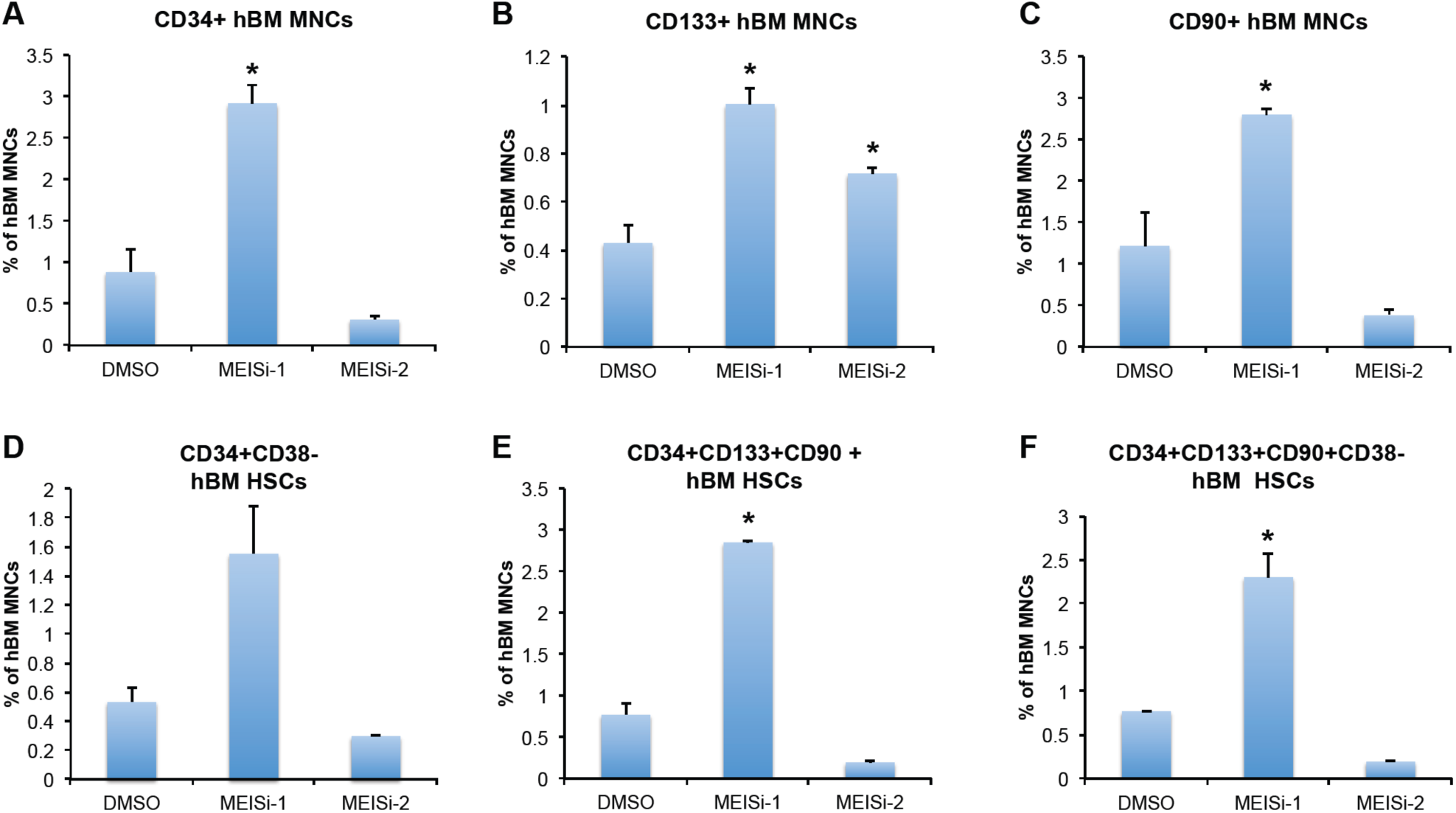
Human bone marrow hematopoietic cell expansion post MEISi treatments. **A)** the percentage of CD34^+^ hBM MNC content, **B)** the percentage of CD133^+^ hBM MNC content, **C)** the percentage of CD90+ hBM MNC content D) the percentage of CD34^+^CD38^-^ hBM MNC content, **E)** the percentage of CD34^+^CD133^+^CD90^+^ hBM MNC content, **F)** the percentage of CD34^+^CD133^+^CD90^+^CD38^-^ hBM MNC post 7 days of treatment with MEISi-1 and MEISi-2 compared to the cells treated with DMSO. n=3, *p <0.05.

### 3.7 MEISi induces HDR/S phase gene expression in hematopoietic cells

One of the key attributes and success in gene editing is efficient HDR/S-phase related gene expression and cell cycle progression. Thus, we determined HDR/S-phase related gene expression profile of MEISi-1 and MEISi-2 treated Lin-cells. We have found that MEISi-1 and MEISi-2 treatments lead to upregulation of *PCNA* gene expression upto 100 fold, and Mcm2 gene expression up to 5 fold (**Figure S9**). In addition, we studied effect of MEISi-1 and MEISi-2 in bone marrow derived or adipose derived mesenchymal stem cells and endothelial cells. To this end, when we treated BM-MSCs, HUVECs or AD-MSCs with MEISi-1 and MEISi-2, we did not see any significant effect in the cell growth post 3 days of treatments (**Figure S10**). These findings suggest that proliferative effect is through upregulation of key S-phase genes and this proliferative effect is unique to hematopoietic cells.

### 3.8 MEISi treated HSCs successfully engraft and repopulate immunodeficient mice

In order to assess the proper expansion of HSCs *ex vivo* with MEISi treatments, we have transplanted HSCs expanded with MEISi treatments into immune deficient animals that lack any functional T or B cells. Procedure involved isolation of CD45.2 HSCs from wild type mice by FACS and expansion with MEISi-1, MEISi-2 or DMSO added HSC medium for 7 days. This was followed by transplantation into CD45.1+ NOD/SCID recipient mice. Then, we have collected blood from recipient mice retroorbitally post 1 and 4 months for assessment of short-term and long-term repopulation, respectively. We have found that MEISi induced HSCs successfully repopulated recipient mice post 1 and 4 months as indicated by presence of CD45.2+ cells in PB (**Figure 6A**). Transplanted CD45.2+ HSCs also successfully differentiated into granulocytes/macrophages, T cells and B cells (**Figure 6B**).

**Figure 6.**
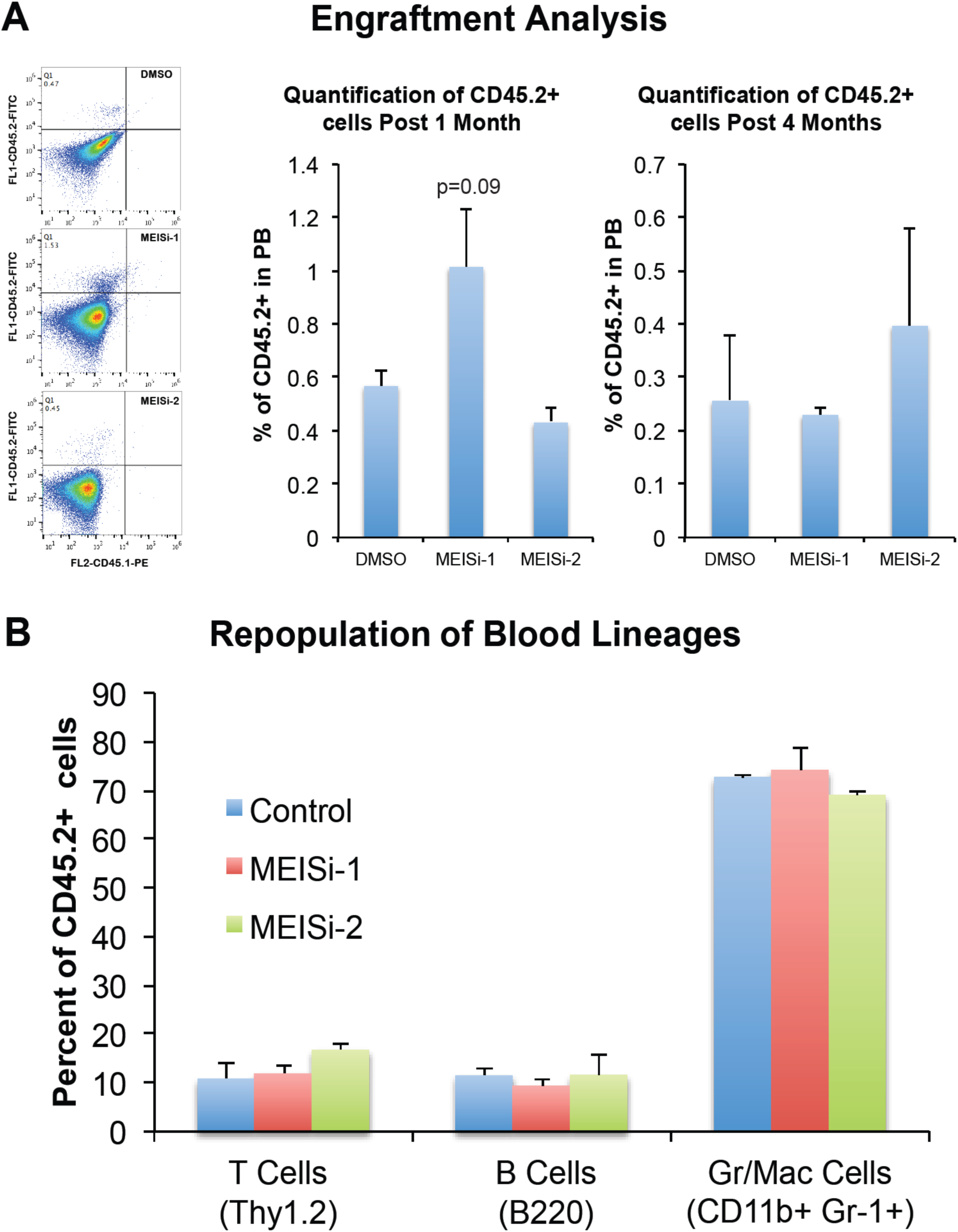
Analysis of engraftment and repopulation post MEISi expanded HSC transplantation. Murine LSK cells expressing CD45.2 allele were treated with effective dose of MEIS inhibitors and expanded for 7 days in HSC medium. These expanded HSCs were injected to NOD/SCID mice expressing CD45.1allele in a retro-orbital way. After 1 and 4 months, peripheral blood analysis was performed. Flow cytometry analysis was done with anti-CD45.2-FITCH and anti-CD45.1-PE. **A) Engraftment analysis.** Engraftment and repopulation of HSCs were quantified based on the percentage of CD45.2^+^CD45.1^-^ cells in the peripheral blood. **B) Repopulation of blood lineages.** Quantification of B cells, granulocytes/macrophages and T cells derived from CD45.2^+^ HSCs in the recipient were determined. n=4.

## Discussion

*In silico*, *in vitro*, *ex vivo* and *in vivo* approaches were employed to develop small-molecule MEIS inhibitors. *In silico* screening of over a million druggable small molecules allowed to identify putative MEISi. MEIS dependent luciferase reporter assays were used to validate *in vitro* efficacy of MEIS inhibitors. Small molecules named MEISi-1 and MEISi-2 demonstrated over 95% inhibition of MEIS luciferase activity. In addition, inhibition of MEIS protein resulted in downregulation of MEIS1 target gene expression in human and animal models *ex vivo* and *in vivo*, respectively. It was found that MEISi-1 and MEISi-2 reduced expression of *Hif-1α, Hif-2α* and *p21* genes in the Meis1 pathway ^5,13,14,16,17^. Intriguingly, MEISi-1 and MEISi-2 demonstrated different dynamics in terms of dosage, target gene modulation and specificity. Both MEISi-1 and MEISi-2 inhibited luciferase reporter at 100 nM, however, MEISi-2 activity seemed to get lower in higher doses. MEISi-2, similarly, did not inhibit PBX1-Luc reporter while MEISi-1 demonstrated inhibition of PBX-Luc reporter at higher doses (10 µM). These findings suggest that MEISi-1 could be a potential and broad PBX1 and maybe TALE family of homeodomain inhibitor at 10 µM and higher concentrations.

Small molecules are desired therapeutic agents that differ in important ways from other types of drug compounds (e.g., proteins, or biologics). Their small size allows them to penetrate inside of the cells and inhibit target moieties. However, small molecules, when used in high doses, demonstrate non-specific inhibition of similar target moieties. Therefore, small molecule inhibitors targeting a specific protein or pathway are expected to have high affinity to target but low affinity to homologous/highly similar proteins. In this sense, we have performed *in silico* screening of MEIS HD inhibitors by comparing to highly similar members of the same TALE family of proteins. Among these proteins, PKNOX1 is also known to have tumor suppressor function and TGIF1 is reported to act as negative regulator in MLL-rearranged AML ^36^. We selected hits that have low affinity towards PKNOX1 HD and TGIF1 HD. Thus, MEIS inhibitors are selected among hits that do not inhibit TGIF1/TGIF2 or PKNOX1. In addition, we eliminated hits that have high affinity to PBX HD to identify highly specific MEIS inhibitors. Furthermore, we have applied another filter to eliminate any potentially cardiotoxic compounds by determining affinity to hERG channel ^24,37^. These findings were further validated by the analysis of *Pbx, Tgif1, Tgif2* and *pKnox* gene expression and their target gene expression profile post MEIS inhibitor treatments. We have found that while we achieve significant downregulation of *Meis1* and *Meis2* expression (**Figure S5**), we did not observe any significant change in expression profile of majority of related TALE family of genes analyzed. We have observed downregulation of Hoxa9 expression to some extent. When MEIS inhibitors are used in cancer studies, downregulation of *Hoxa9* could be beneficial because MEIS1 is known to cooperate with HOXA9 in carcinogenesis.

Studies have shown that *Meis1* has fundamental roles in heart regeneration, carcinogenesis and stem cell function. Previously, we have demonstrated that MEIS1 limits neonatal cardiac regeneration^5^. Knocking out *Meis1* in cardiomyocyte specific manner leads to improved cardiac function by extension of cardiac regeneration window in neonatal mouse. This suggests that MEIS inhibitors could be utilized in the activation of cardiac regeneration post cardiac injuries.

Meis1 was first discovered in cancer as a viral integration site. Since then, over hundred different studies demonstrated a correlation between tumorigenesis and dysregulation of Meis1 expression. Meis1 has been shown to be over expressed in a number of cancers and been indicated with its oncogenic potential ^38–40^. *Meis1* transcriptionally regulates the expression of hypoxic tumor markers, *Hif-1α* and *Hif-2α*. *Hif-1α* and *Hif-2α* are involved in the induction of glycolysis and scavenging of reactive oxygen species. MEIS1 along with other homeobox proteins often found to be driving hematopoietic transformation in MLL-rearranged leukemia and described as an oncogene. Intriguingly, a high level of Meis1 expression was found to be associated with resistance to conventional chemotherapies. Although correlations have been made regarding *Meis1* and tumorigenesis, the molecular mechanism behind it remains undetermined (reviewed in^5^). Newly developed MEIS inhibitors could be used to address questions regarding the involvement of MEIS proteins and underlying pathways in the tumorigenesis, and maybe used to develop a novel anti-cancer therapies for various cancer types.

HSCs are recognized by their self-renewal and differentiation potential to all blood cell types. They are responsible for generation of billions of mature blood lineages throughout life of an adult ^5,34^. HSC transplantation has been used for decades in the treatment of leukemia, lymphoma, some solid cancers, autoimmune diseases and inherited diseases like sickle cell anemia ^41–45^. Studies showed that gene edited HSCs could be used for treatment of genetic diseases ^41–43,46–49^. However, these procedures are still limited due to inefficient *ex vivo* expansion of gene edited HSCs.

We previously showed that HSC specific deletion of *Meis1* results in expansion of HSC pool *in vivo* ^13^. HSCs are could be expanded by treatment of cytokines and growth factors such as TPO, Flt-3l, SCF albeit with tendency to differentiate when they are kept in culture for long periods. Thus, we have assessed the effect of MEIS inhibitors to shorten HSC expansion process by targeting MEIS related quiescence. We have found that MEIS inhibitors effectively enable functional increase of the both mouse and human HSCs in the presence of TPO, Flt-3l, SCF. This was evident from the increasing number of CFU-GEMM colonies, and induction of HSC cells analyzed by flow cytometer. It was also demonstrated that *in vivo* application of MEISi-1 and MEISi-2 successfully increased HSC number in mouse bone marrow (LSKCD34low cells). These studies suggest that MEIS inhibitors can be adapted to alternative stem cell harvesting and induction of HSCs expansion technologies. These studies highlight emerging use of MEIS inhibitors in the development of new therapeutic approaches in the treatment of cardiac injuries, hematopoiesis issues, bone marrow transplantations, and cancer.

## Acknowledgments

We like to thank to the support by The Scientific and Technological Research Council of Turkey (TÜBİTAK) ARDEB 3501 [#215Z071] program and Turkish Hematology Association 2016 Research Project Award. FK is supported by funds provided by the European Commission Co-Funded Brain Circulation Scheme by The Marie Curie Action COFUND of the 7th. Framework Programme (FP7) (115C039), The Scientific and Technological Research Council of Turkey (TÜBİTAK) [grant numbers 115S185, 215Z069, 215Z071,and 216S317], The Science Academy Young Scientist Award Program (BAGEP-2015, Turkey), The International Centre for Genetic Engineering and Biotechnology – ICGEB 2015 Early Career Return Grant [grant number CRP/TUR15-02_EC], Medicine for Malaria Venture MMV Pathogenbox Award (Bill and Melinda Gates Foundation), Gilead Sciences International Hematology & Oncology program, Gilead ile Hayat Bulan Fikirler, ERA-Net CVD program. EA is supported by TÜBİTAK BIDEB 2211-A program. MU has been supported by TÜBİTAK 215Z071 and 118S540. We like to thank Enes & Kemal for their help in establishment of small molecules targeting homeobox library of proteins and facilitating the re-docking studies into other TALE family members. We are also thankful to Mine Yarim Yüksel and Ece Gürdal from Faculty of Pharmacy, Yeditepe University for her technical advises in molecular dockings and interactions. We are thankful to Serli Canikyan from Onkim Stem Cell Technologies for her technical advises in UCB stem cell collection.

## Author Contributions

F.K designed the all experiments and performed majority of *in silico* studies, wrote the article. R.D.T. performed luciferase assays, RT-PCR, flow cytometry analysis and *in vivo* studies. L.Y.A. and G.S.A. performed and contributed to RT-PCR. E.A., P.S., N.M., Z.G. performed and contributed to human HSC studies. E.A. performed AD-MSC studies. D.Y. and B.M.K performed HUVEC studies. L.Y.A. and M.A. performed and contributed to BM-MSC studies. E.C.T. contributed to flow cytometry analysis of apoptosis and cell cycle analysis. S.D. performed cardiotoxicity predictions. M.U. performed additional luciferase assays. All authors contributed to data analysis and reviewed the manuscript.

## Figures and Figure Legends

**Figure S1 (related to Figure 1).**
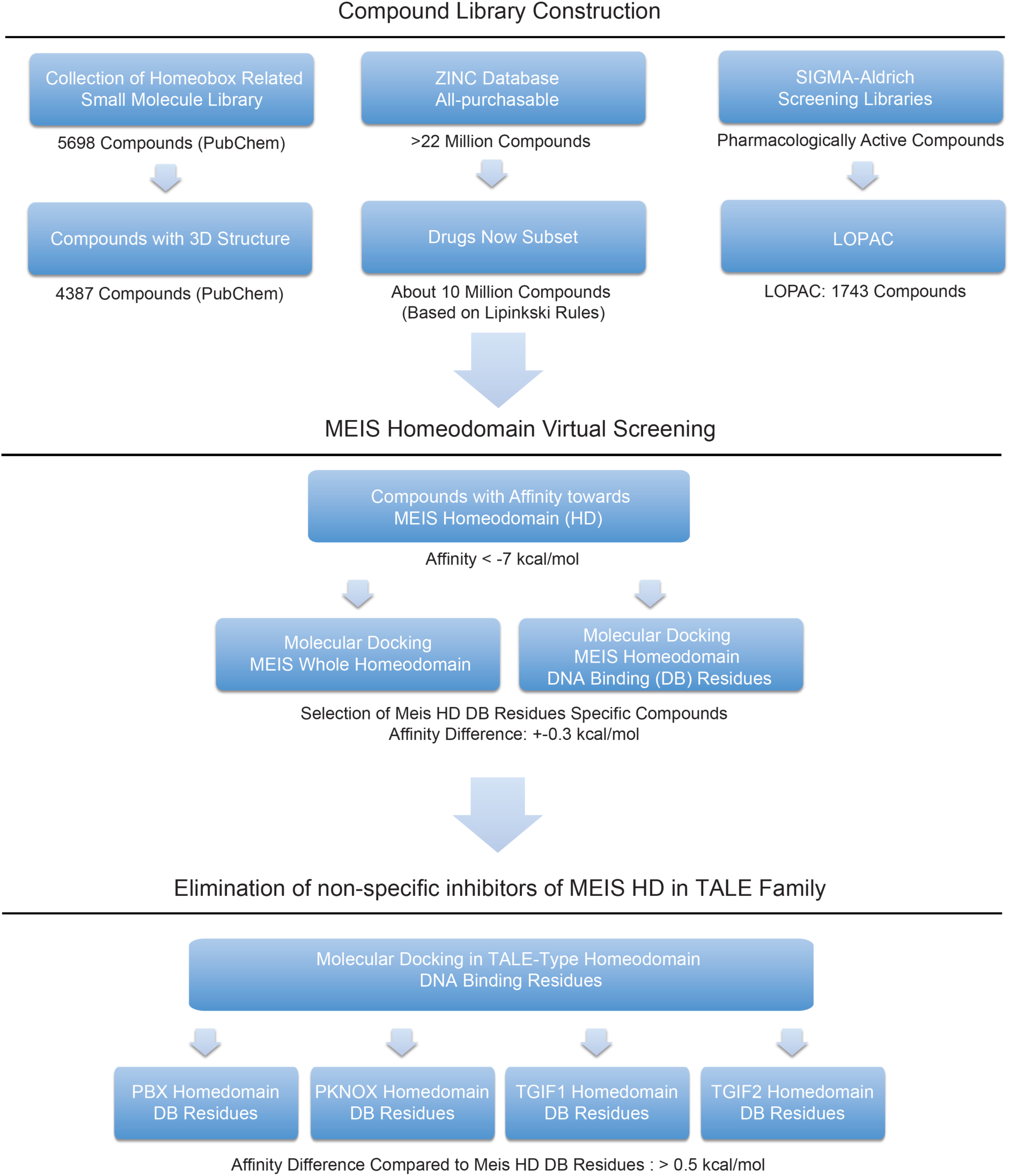
Virtual Screening Strategy. Virtual screening strategy is a tree step process. Initially, *in silico* screening library were generated by compiling SDF files from ZINC drugs-now subset, Sigma LOPAC1280 and curated inhibitors of homeobox family of proteins from PubChem. In the second step, small molecules were docked into whole MEIS homeodomain as well as DNA interacting-highly conserved residues. Final step includes affinity calculation of selected hits against other TALE family proteins to eliminate non-specific hits.

**Figure S2 (related to the Figure 1).**
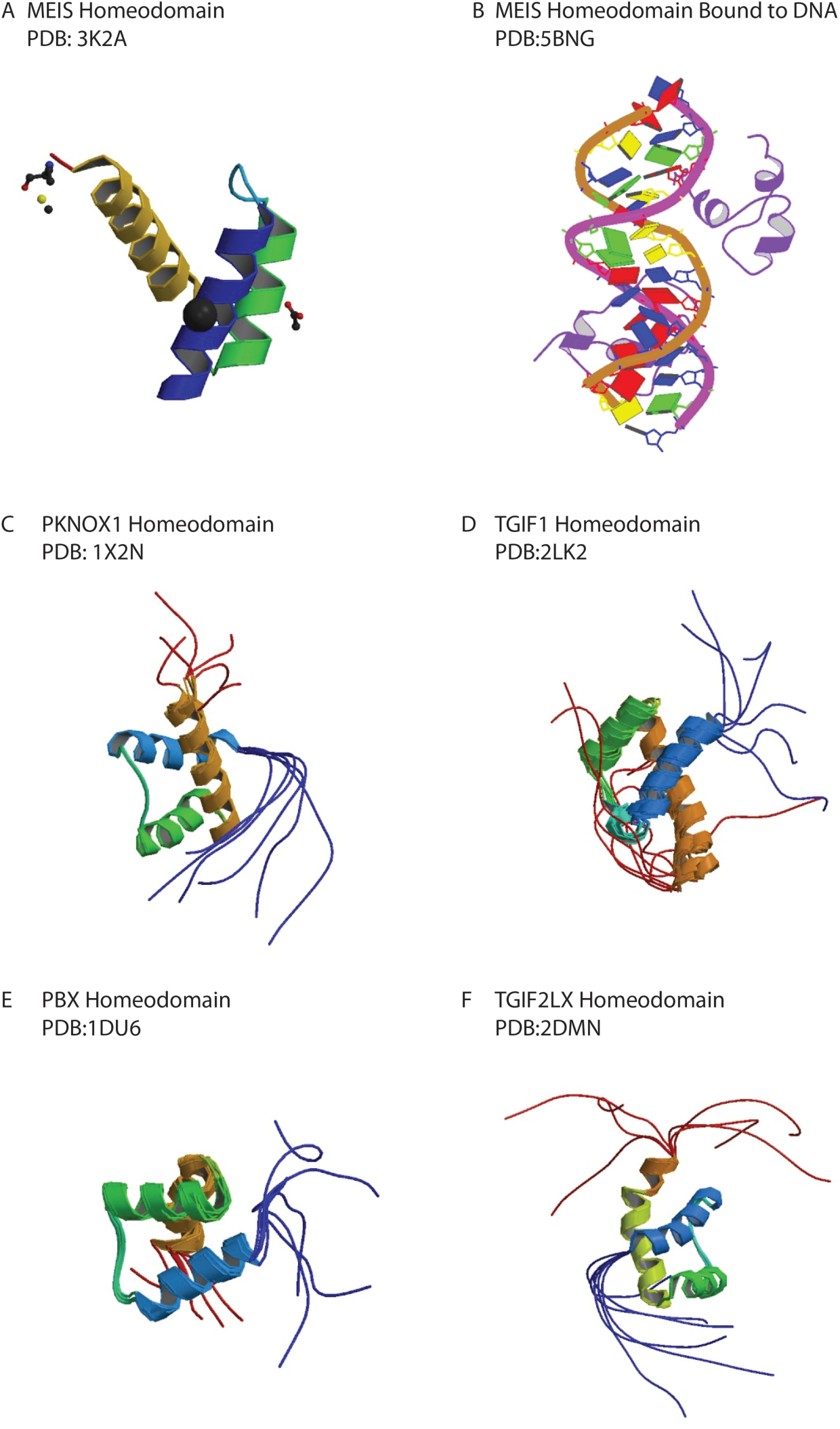
Crystal Structure of TALE-type homeodomains. TALE family proteins with known homeodomain crystal structure have been obtained from PDB and docked with MEIS small molecule hits. Gibbon diagrams for **A)** MEIS HD, **B)** MEIS HD with DNA, **C)** PKNOX HD, **D)** TGIF1 HD, **E)** PBX HD, **F)** TGIF2LX HD are shown. Grid boxes for docking were generated around DNA binding helices

**Figure S3 (related to Figure 2).**
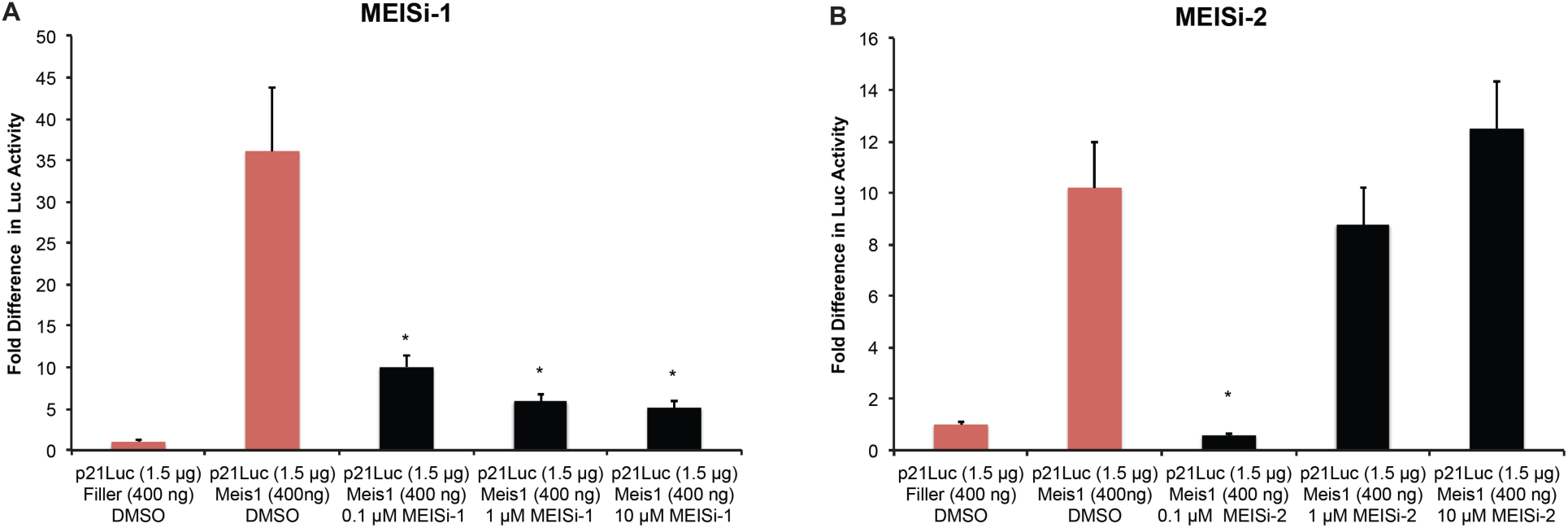
Dose dependent effect of MEISi-1 and MEISi-2 in MEIS luciferase reporter activity. Luciferase reporter assays demonstrate the dose dependent effect of **A)** MEISi-1 and **B)** MEISi-2 compared to DMSO control. n=3, *p<0.05.

**Figure S4 (related to Figure 2).**
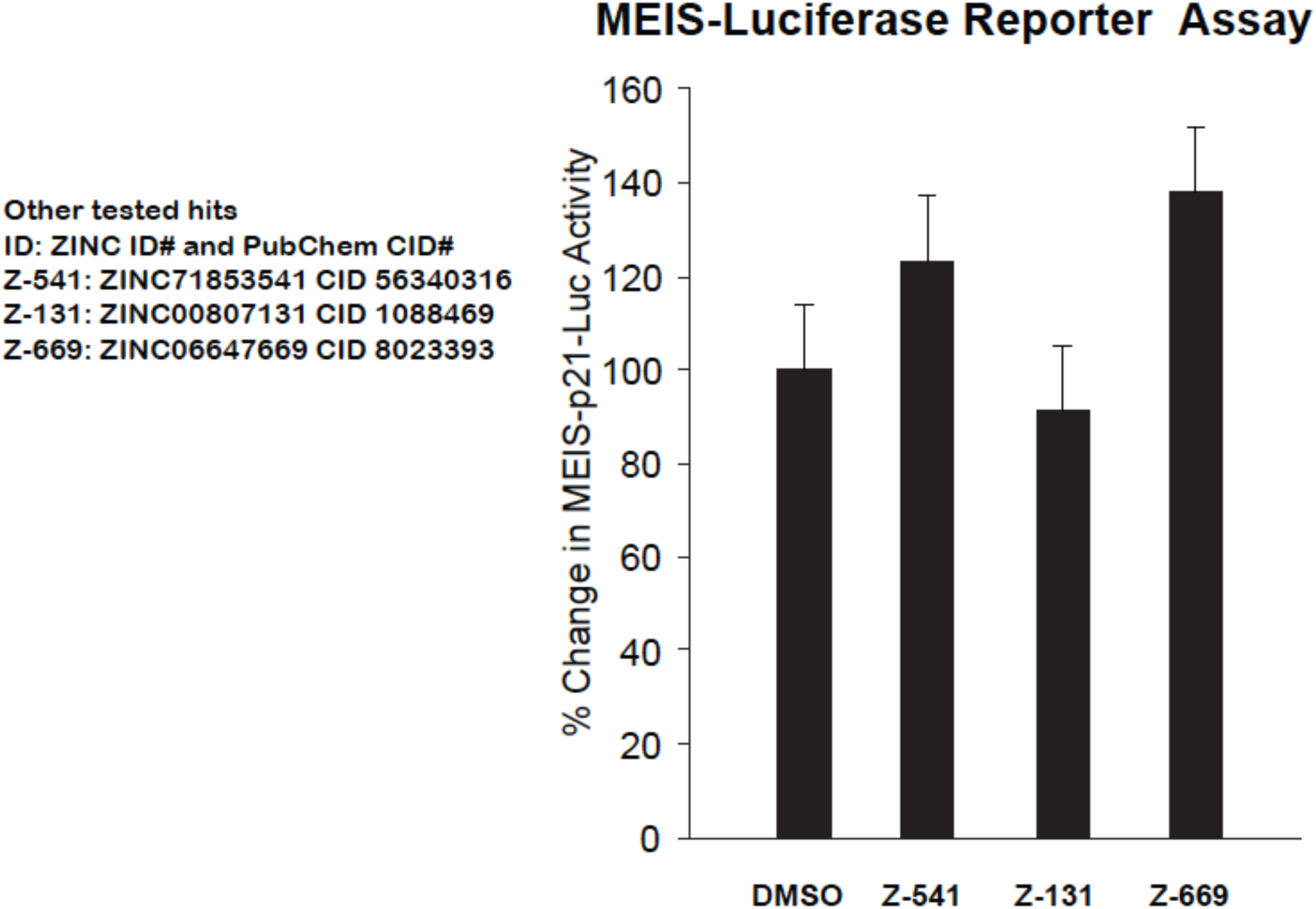
Other tested MEIS hits with similar structure. Analysis of other hits similar to MEISi-1 and MEISi-2 in MEIS luciferase reporter assay.

**Figure S5 (related to Figure 2).**
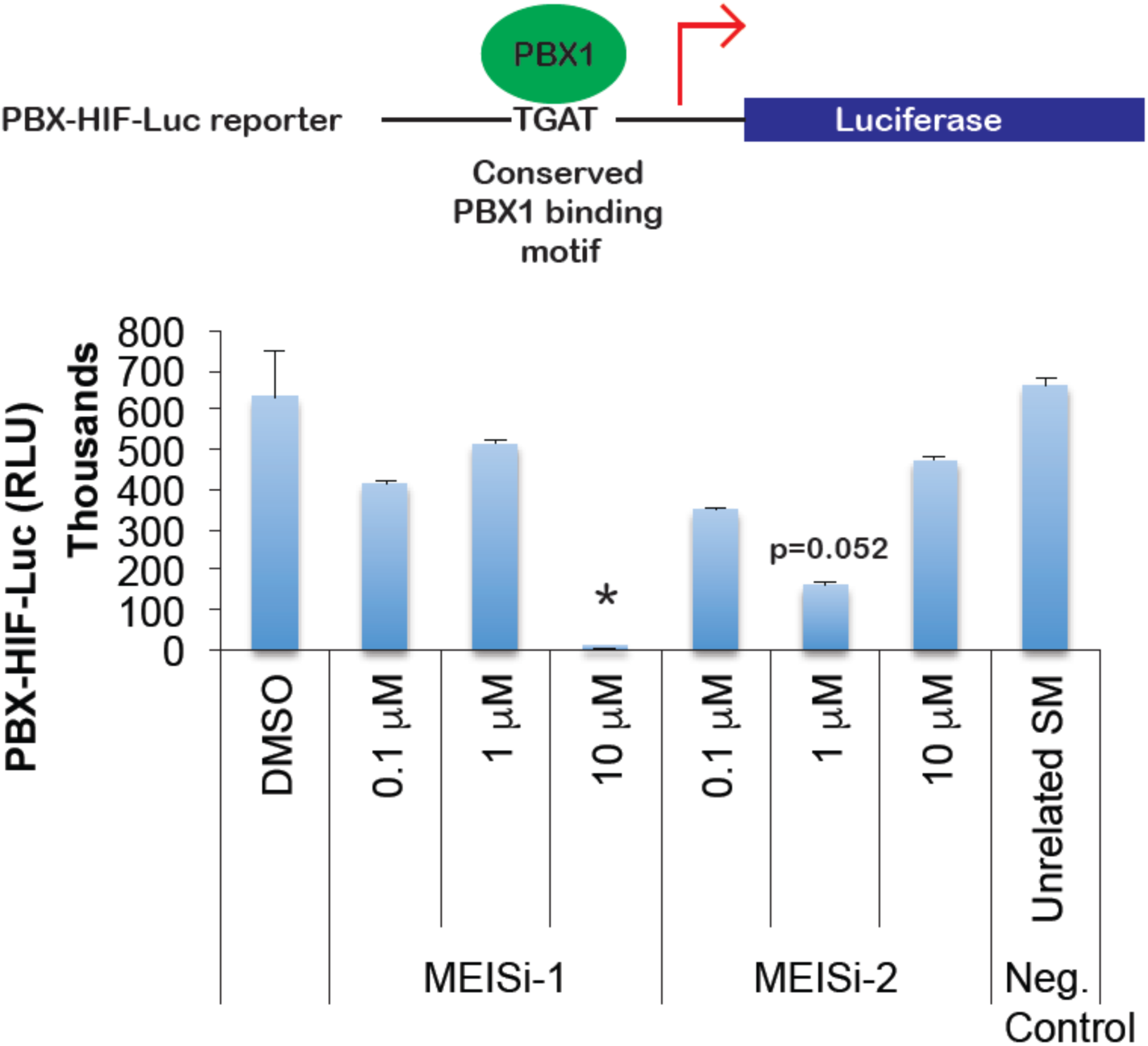
MEIS inhibitors tested in PBX-Luc Reporter. Analysis of other MEISi-1 and MEISi-2 in PBX-luc-reporter assay. n=3, *p<0.05.

**Figure S6 (related to Figure 2).**
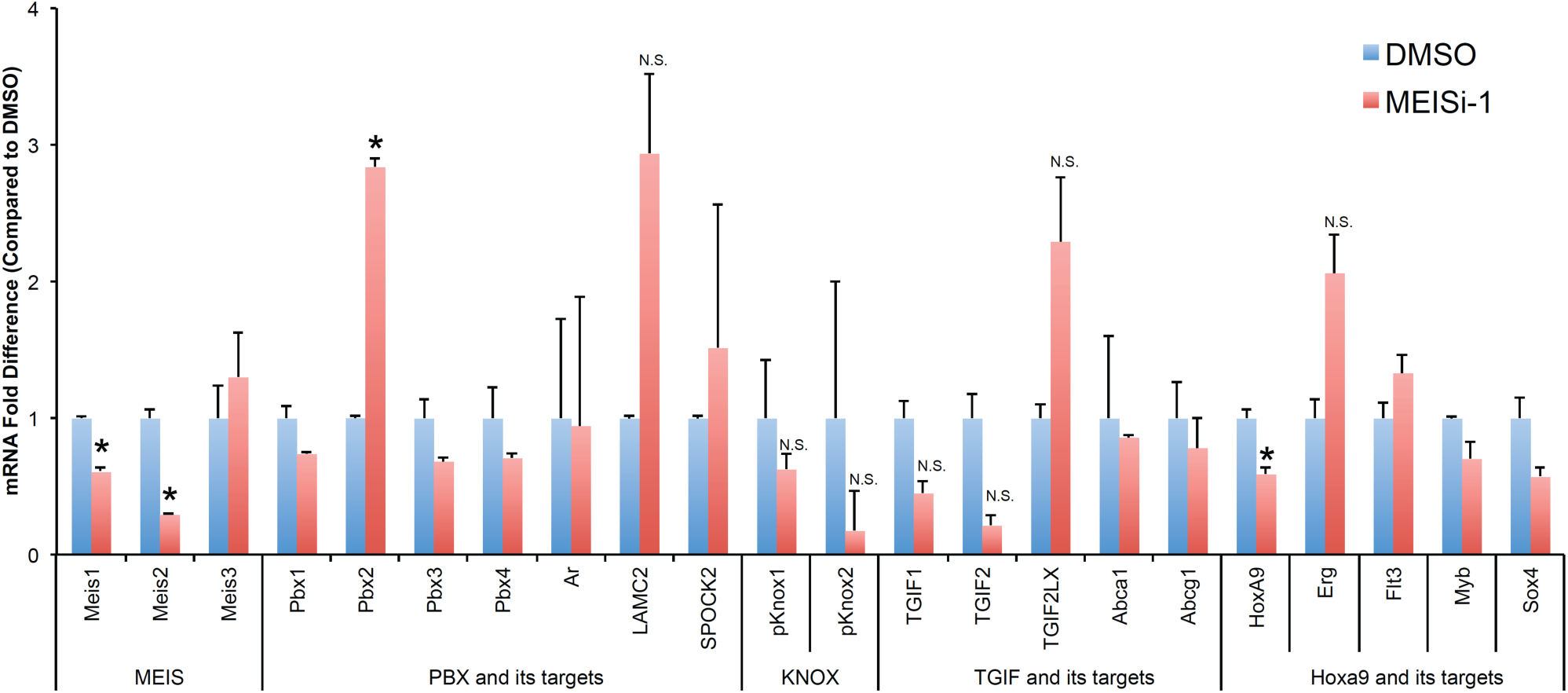
Gene expression analysis of TALE family genes and their targets. We have determined gene expression analysis post MEISi-1 to determine how MEIS inhibitors affect expression of Meis1, Meis1 related, other TALE family genes, and more importantly their target genes. n=3, *p<0.05, N.S.=Not significant.

**Figure S7 (related to Figure 2).**
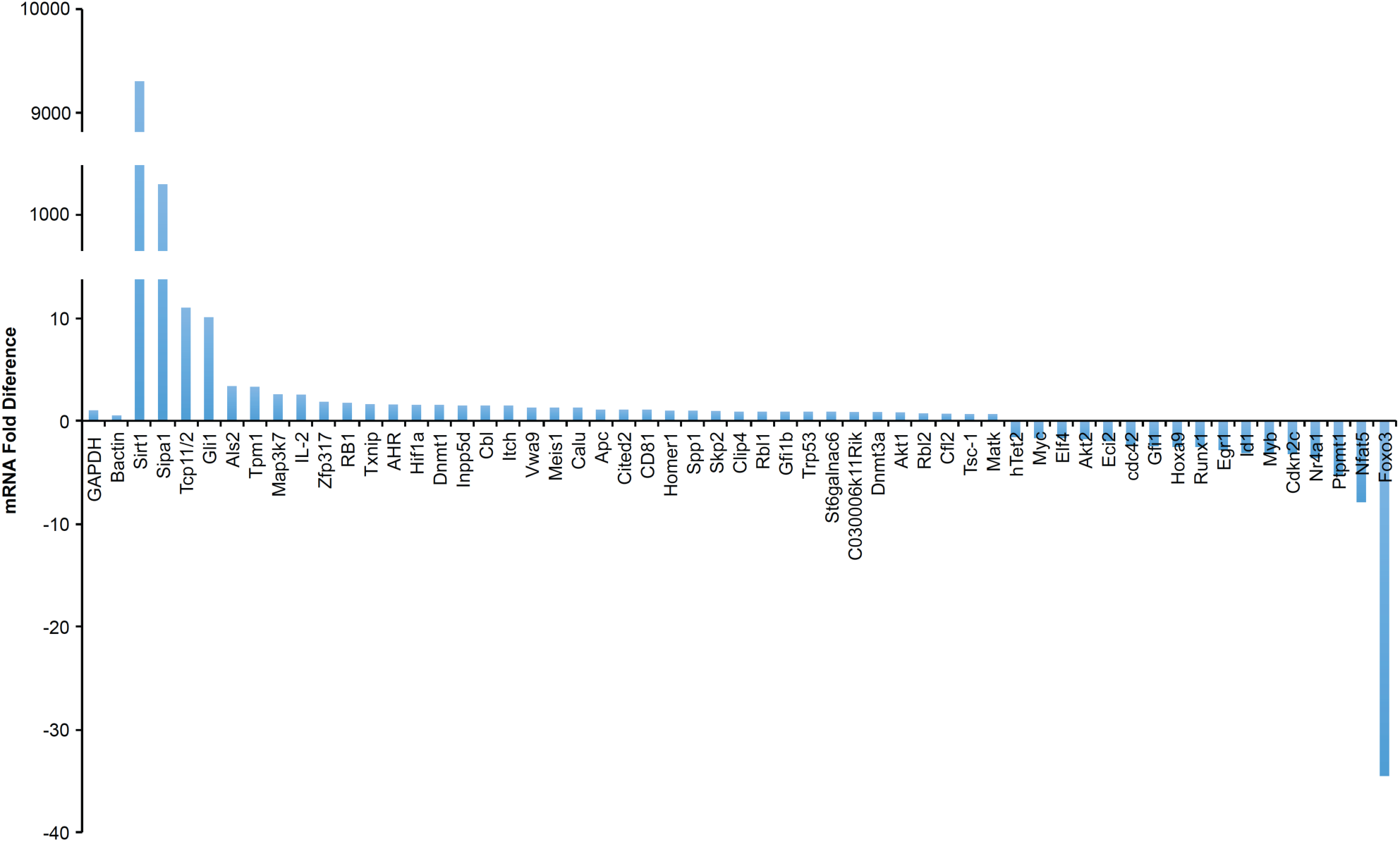
Analysis of HSC gene pool by PCR array post MEISi-1 treatment in Lin-Cells. We have determined gene expression analysis post MEISi-1 to determine how MEIS inhibitors affect expression of HSC related genes.

**Figure S8 (related to Figure 4).**
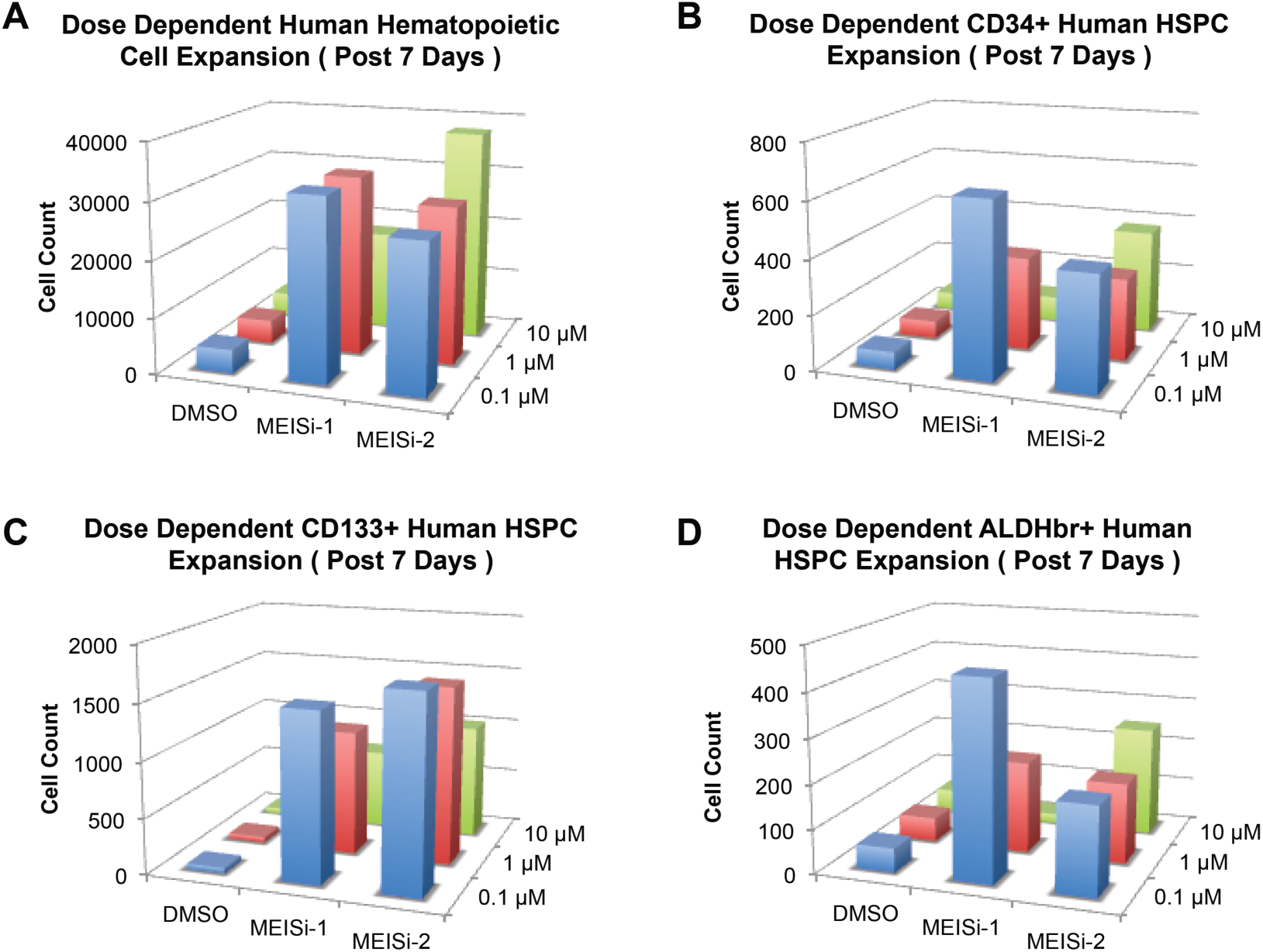
Human UCB hematopoietic cell expansion post MEISi treatments. Human UCB MNCs were treated with three different doses of MEISi-1 and MEISi-2. **A)** Human hematopoietic cell count, **B)** CD34^+^ human HSPC, **C)** CD133^+^ human HSPC count, **D)** ALDH^br^ human HSPC counts were determined at given concantrations.

**Figure S9.**
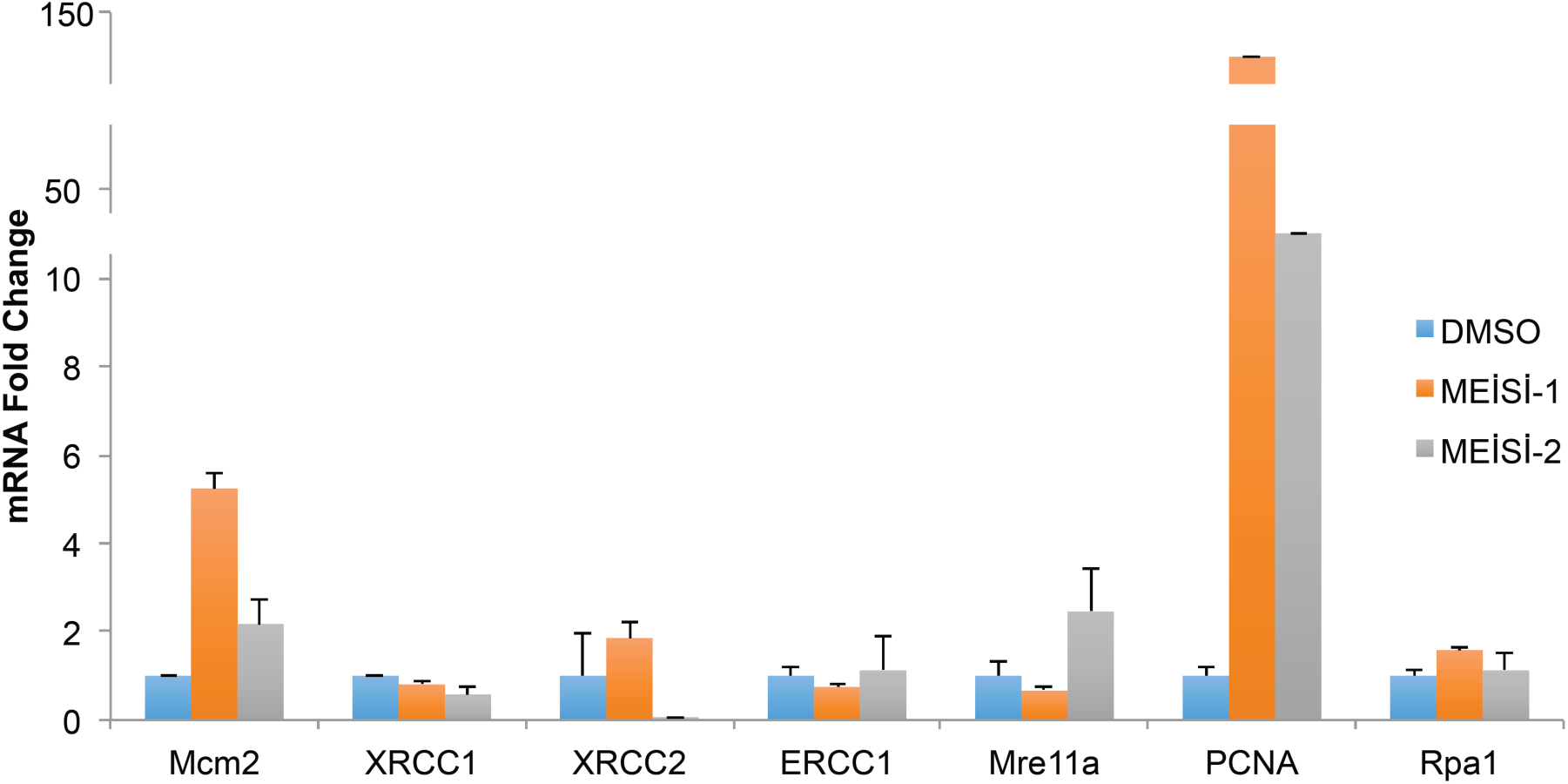
Effect of MEISi treatments in the expression of HDR related genes. Lin-cells were treated *in vitro* with MEIS inhibitors and collected RNA post 3 days of treatment for analysis of gene expression. n=3.

**Figure S10.**
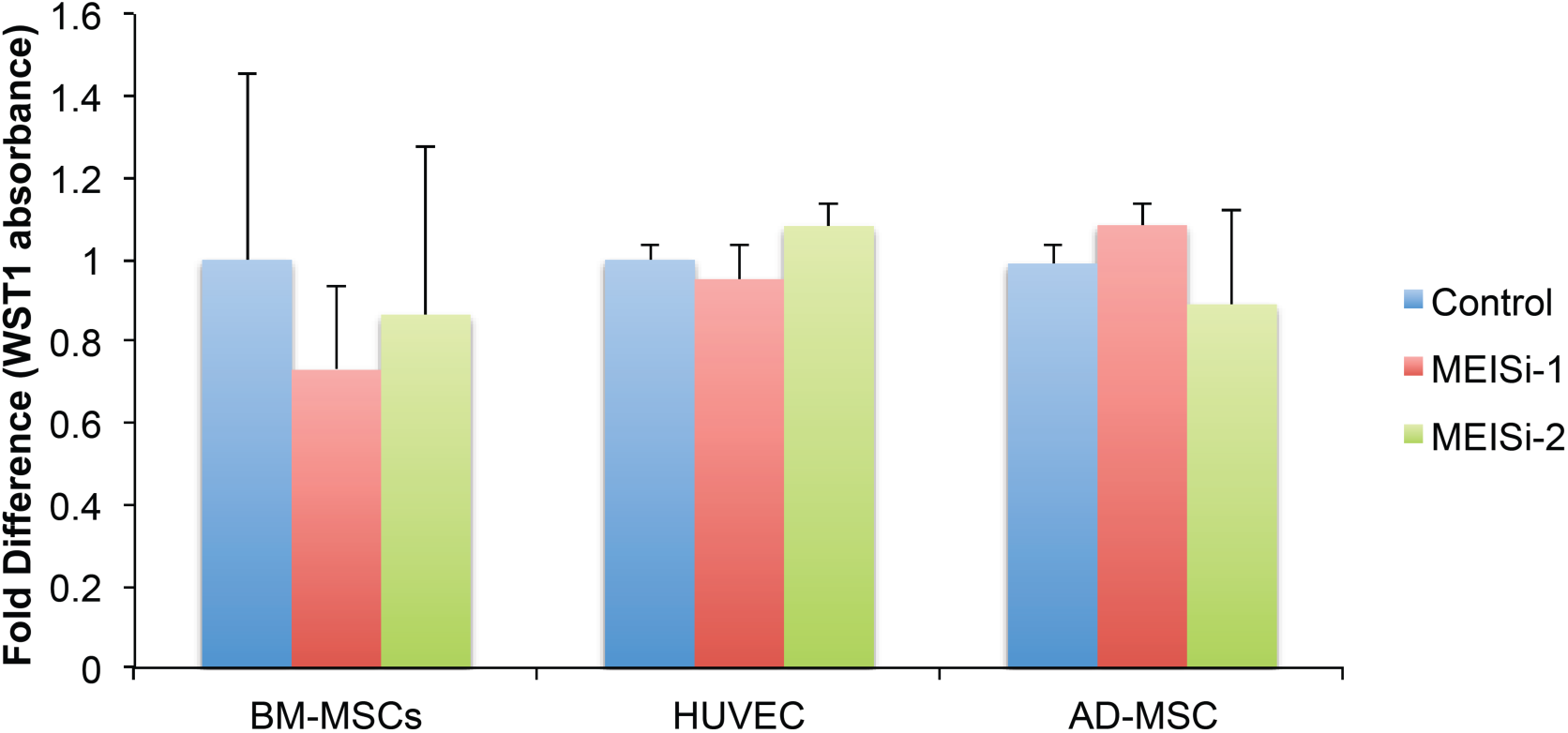
Effect of MEISi treatments in BM-MSC, HUVEC and AD-MSC proliferation post 4 days. BM-MSCs, HUVECs, and AD-MSCs were treated with 1 µM of MEISi-1, MEISi-2 inhibitors and DMSO (control, 0.5%). Fold difference in WST1 absorbance was determined after 72 hours. n=3.

## Supplementary Files

**Supplementary File 1 (related to Figure 1)**: *in silico* screening of ZINC drugs-now small molecule database & MEIS HD (whole) vs MEIS DB (DNA binding residues) affinity calculations.

**Supplementary File 2 (related to Figure 1): Enrichment analysis.**

**Supplementary File 3 (related to Figure 1 and Table 5):** Selected hit small molecules with no predicted cardiotoxicity. Hit molecules were docked with Glide/XP into hERG. Affinities predicted to be <-7.8 kcal/mol (Maestro Glide) or <-7.4 kcal/mol (AutoDockVina) are considered potentially cardiotoxic.

## Data availability

Additional data available upon request.

